# Comparative investigations of cellular dynamics in the development of medusae (Cnidaria: Medusozoa)

**DOI:** 10.1101/2024.04.28.591574

**Authors:** Matthew Travert, Kent Winata, Paulyn Cartwright

## Abstract

Medusozoans are characterized by the presence of a medusa (jellyfish) stage as part of the life cycle. Despite being a prominent trait in medusozoan evolution, the mechanisms underlying the emergence of this life cycle stage are poorly understood. Characterizing cell proliferation, cell migration and programmed cell death in several disparate species, we found that the development of the medusa displays distinct cellular mechanisms between the scyphozoan and hydrozoan lineages. Using Edu labeling, hydroxyurea treatments and cell tracing we found that in hydrozoans, the development of the medusa relies on extensive cell migration and a distinct pattern of cell proliferation. Using TUNEL assays and pan-caspase inhibitor treatments, we found that in all surveyed hydrozoans apoptosis does not play a role in the development or release of medusa. By contrast, the surveyed scyphozoan medusae undergo extensive apoptosis, and subsequent cell proliferation is required for the formation of the medusa and many of their adult structures. Our results suggest that the difference in medusa developmental trajectories between scyphozoans and hydrozoans stems from distinct cellular mechanisms. Characterization of the medusa stage at different levels of biological organization is thus required to investigate the origin of this complex life cycle stage.

## Introduction

The medusa, commonly referred to as jellyfish, is a key component in the life cycle of medusozoans. Medusa is typically used to refer to the pelagic, free-living, sexually reproductive stage in the life cycle of medusozoans, and is thus effectively defined in a functional paradigm, without reference to specific morphological criteria. The group Medusozoa was originally proposed by Petersen (1979) and was based on the grouping of cnidarians possessing a medusa stage in their life cycle. Study of mitochondrial genome structure further supported the clade Medusozoa (Bridge et al., 1995) that was ultimately solidified by multiple phylogenetic analyses (Zapata et al., 2015, Collins et al., 2006, Simion et al., 2017). The subphylum Medusozoa (Petersen, 1979) is divided into two groups, Hydrozoa that possesses a hydromedusa and Acraspeda (Gegenbaur, 1856), composed of Scyphozoa (true jellyfish), Staurozoa (stalked jellyfish) and Cubozoa (box jellyfish). Stemming from this functional definition, the single origin of the medusa is often assumed, being found across Medusozoa. Nonetheless, the medusa represents a multifaceted trait with a complex evolutionary history. Within medusozoans, the medusa exhibits tremendous variation in its morphoanatomy and development, consistent with their life cycles.

The typical medusozoan life cycle is composed of a planula larva, a polyp, and a medusa stage. Throughout medusozoan evolution, multiple instances of alteration of the life cycle have occurred and loss of the planula or polyp, although rare, are found in both Acraspeda (Jarms et al., 1999) and Hydrozoa (Petersen, 1990 and Bentlage et al., 2018). However, loss, reduction and putative regains of the medusa stage are frequently found across hydrozoans and the loss or reduction of the medusa stage has been inferred to have occurred several times independently (Cartwright and Nawrocki, 2010, Leclere et al., 2009).

Given the ancient and complex evolutionary history of medusae, a thorough comparative analysis between the different medusozoan lineages is required in order to better understand the ancestral traits present at the origin of Medusozoa.

### An overview of medusozoan anatomy across lineages

The primary differences in morphoanatomy are found between the two main medusozoan groups, Hydrozoa and Acraspeda (Cubozoa, Scyphozoa, Staurozoa), and relate to their unique structure-function relationships. A medusa is typically the free-living swimming stage that aids in dispersal. In that regard, the hydrozoan medusa (hydromedusa) and the acraspedote medusa possess striated muscles, a unique feature within cnidarians associated with active swimming (Boelsterli, 1977). This subumbrellar circular musculature, often referred to as the coronal muscle, is found in the same location in both the hydromedusa and the acraspedote medusa (Lesh-Laurie and Suchy, 1991, Seipel and Scmid,2005). The hydromedusa differs from acraspedote medusae in the presence of gap-junctions that regulate the coronal muscle (Satterlie, 2008). Additionally, the hydrozoan coronal muscle expands in a hydrozoan-specific membranous structure at the base of the bell called the velum (Carré and Carré, 1994). In both types of medusae, the striated muscle is ectodermal in origin, although in most hydrozoans the striated muscle develops from an ectodermally derived primordium called the entocodon (Kuhn, 1910).

The locomotion of the medusa is tightly regulated by the nervous system but in this aspect, the hydromedusa and the acraspedote medusa greatly differ. The hydromedusa possesses two nerve rings, an inner ring connecting the velum and an outer ring running through the bell margin and connecting the sensory system (Thomas & Edwards 1991) in addition to a diffuse nerve net in the manubrium, the canals, and the subumbrella (Koizumi et al., 2014, Groger and Schmid, 2000). The acraspedote medusa is characterized by a diffuse nervous system present throughout the epidermis (Horridge 1956; Lesh-Laurie & Suchy 1991). The nervous system exhibits greater densification and integration in the marginal sensory organs called rhopalia/rhopalioid (Nakanishi 2009, Westlake and Page, 2016). The rhopalium is a tentacle-derived hollow structure bearing statocysts and photoreceptors and is found only in Acraspeda (Berrill,1949). Cubozoans also have a single nerve ring in the bell in cubozoans (Garm et al., 2006, Nielsen et al., 2021).

In connection to the nervous system, various sense organs are found across medusozoan lineages, predominantly functioning in balance (statocysts) and vision (eyes). The eyes of the medusa are found as either eyespots, pigment cups or lensed eyes and have been shown to be convergent (Picciani et al., 2018). Statocysts are found in the Acraspeda groups Cubozoa and Scyphozoa and in two distantly related lineages of Hydrozoa, Leptothecata and Trachylinae (Bouillon 1985, 1994b). The statocysts of cubomedusae, scyphomedusae and trachyline hydromedusae are gastrodermal in origin and share the same statolith mineral composition (Mayer 1910). By contrast, the statocysts in the leptothecate hydromedusae are of epidermal origin with a unique mineral composition and are inferred to have evolved independently in Leptothecata (Chapman 1985; Lesh-Laurie & Suchy 1991). These differences between lineages, as well as the absence of statocysts in Staurozoa and many hydrozoans, suggest possible convergent evolution of this trait. Statocysts may have been present in the last common ancestor of medusozoans and lost in Staurozoa and non-trachyline hydrozoans (Hydroidolina) and re-evolved in Leptothecata. Alternatively, statocysts could have evolved independently in Rhopaliophora (Scyphozoa and Cubozoa), Trachylina and Leptothecata.

In most cases the gastrovascular system of the medusa is composed of a gastric cavity vascularizing the bell via radial canals connected to a ring canal running through the bell margin. A complete gastrovascular system is found in all fully developed hydromedusae, but Narcomedusae lack radial canals, and exhibit a system analogous to a ring canal (peripheral canal, Bouillon 1994a; Carré and Carré 1994; Bouillon 1978). In acraspedote medusae the radial canals are only found in scyphomedusae, and the ring canal is found only in a subset of scyphomedusae (Thiel 1966). The gastric cavity of the acraspedote medusa is septated and characterized by the presence of gastric filaments.

Often associated with the gastrovascular system are the gonads. In Acraspeda the gonad is gastrodermal in origin and location and is found in the form of gastrodermal folding (Adonin et al., 2012, Garcia-Rodriguez et al.,2018, Berrill, 1963). In hydrozoans the gonad is loosely defined and constitutes a regionalized and functionally specialized structure composed of an outer and inner epithelial layer where the storage and maturation of the gametes take place (see Bouillon et al 2006). In hydrozoans, gonads may be located on the manubrium or the radial canals and the position is lineage specific (Bouillon 1985; Bouillon 1994b; Mayer, 1910).

Overall, characterization of the traits that constitute the medusa stage leaves substantial ambiguity regarding the ancestral condition within Medusozoa. The morphoanatomy and distribution of traits in extant medusae is consistent with the last common ancestor of Medusozoa exhibiting a manubrium, coronal and striated muscles, a diffuse nervous system in the subumbrella and marginal tentacles without tentacle bulbs. The various remaining structures either cannot be unambiguously inferred (radial canals, statocyst, nerve ring, position, and origin of the gonad) or are inferred to evolve independently in separate lineages (eyes, ring canal).

### Acraspeda and Hydrozoa medusae exhibit drastically different developmental patterns

Typical medusa development occurs from the polyp. Hydromedusa development unfolds through lateral budding, initiated by the out-pocketing of the polyp epithelia, culminating in the formation of a fully developed medusa. The invading gastrodermis ramifies into radial canals that extend to the bell margin, where subsequent radial expansion leads to the formation of the ring canal (Boelsterli, 1977). The development of the hydromedusa is canonically characterized by the formation of the entocodon, an interstitial cell mass derived from the ectoderm, that gives rise to striated muscle (Berrill 1950). Characterization of the development of the nervous system shows that despite being transiently connected to the polyp, the nervous system that characterizes the hydromedusa develops *de novo* (Groger and Schmid, 2000).

Not all hydromedusae develop from a polyp. In some hydrozoan species, the medusa may asexually develop from a medusa. This type of budding may take place in regions where gonads are typically found (manubrium, radial canals), but can also arise from somatic tissue, such as the exumbrella (Berrill, 1950) or more rarely the tentacle bulb, such as seen in *Niobia dendrotentaculata* (Mayer, 1910). The gonad and the tentacle bulb are known niches of adult stem cells (i-cells, Seipel et al., 2004, Chari et al., 2021).

In some species, the hydromedusa develops directly from the planula larva. In these cases, the polyp stage is absent, and an actinula or actinuloid stage may be found. In hydrozoans, this type of direct development is rare and found exclusively in the Narcomedusae and Trachymedusae lineages (Marques and Collins, 2004 and Bentlage et al., 2018).

In Acraspeda, although the development of the medusa varies between the three main lineages, all develop through a transformation from a pre-existing oral-aboral axis, which is typically the oral end of the polyp, but can also be from the planula larva (see below). This is in contrast to metagenetic hydrozoans, in which medusae bud laterally from the polyp. Canonically, scyphomedusa development is carried out by a process of transverse fission, known as strobilation, of a transformed oral/apical end of the polyp. During strobilation the polyp tentacles regress, some of which will form the rhopalia. Strobilation can be monodisk (apical transformation into a single juvenile medusa called an ephyra), such as seen in *Sanderia malayensis,* or polydisk (apico-basal segmentation into disks giving rise to several ephyrae) such as *seen Aurelia coerulea*, the latter is inferred to be ancestral for Scyphozoa (Helm, 2018), whereas monodisk strobilation being only found in Kolpophorae and in some Pelagiidae and Cyaneidae (e.g *Sanderia malayensis* and *Cyanea nozakii*). Previous studies on two scyphomedusae have shown that muscle develops *de novo* in the ephyra and does not require the presence of the polyp muscle cords (Helm et al., 2015). All swimming acraspedote medusae possess a coronal muscle, while the polyp stage musculature mostly consists of adradial epidermal muscle cords (Thiel 1966), suggesting that the coronal muscle in Staurozoa and Cubozoa might develop *de novo* as well. However myogenesis during medusa development in these two lineages remains to be characterized. Little is known about the development of the nervous system during strobilation, however, the pulsating behavior of developing ephyrae in the strobila of *Aurelia aurita,* observed by Horridge (1956) and Schwab (1977a,) suggests that their nervous system is both autonomous and integrated throughout the strobila. Whether the released ephyra retains parts of the strobila nervous system remains unknown.

Although modifications of the scyphozoan life cycle are not as varied as in hydrozoans, direct development has evolved independently at least twice within this group. In *Pelagia noctiluca* (Rottini Sandrini and Avian, 1983), the planula directly develops into an ephyra, while the coronate *Periphylla periphylla* develops directly into a medusa following gastrulation (Jarms et al., 1999). In both species the oral pole of the gastrula/larva develops into the mouth of the medusa. In those species, the morphoanatomy of the adult medusa shows no outstanding variation compared to their close metagenetic relatives. This suggests that despite strikingly different developmental trajectories, scyphomedusa development occurs through transformation of the oral-aboral axis regardless of whether it arises from the polyp or the larva. Less is known of stauromedusa development, but studies have indicated that it is consistent with oral transformation of the polyp. Following the development of additional adradial tentacles, the tentacles of the polyp transform into clusters of capitate tentacles. Later, the gastric filaments and the gonads develop in the gastroderm (Kikinger and Salvini-Plawen, 1995). Thus, the stauromedusa is entirely derived from the polyp. The neuromuscular development during medusa development remains unknown. Similar to some other members of Acraspeda, the development of the cubomedusa occurs through transformation of the oral end of the polyp (Stangl et al. 2002), exhibiting processes ranging from monodisk strobilation (Toshino et al., 2015) to transformation of the entire polyp into a free-living medusa (Straehler-Pohl and Jarms, 2005).

Although there are some elements of medusozoan morphoanatomy that are shared across the clade, there are striking differences in other traits as well as in their overall developmental trajectories. This creates a paradox where some traits show a discrepancy in inferred common ancestry in these different levels of biological organization (development and adult form). The sister-group relationship between Acraspeda and Hydrozoa and superficial similarities observed in the medusa could indicate deep homology of a medusa-like form with subsequent divergence of developmental processes. By contrast, the pattern also supports parallel homoplasy, where the utilization of dissimilar developmental processes led to the parallel evolution of the medusa form in the two main medusozoan lineages.

Disentangling which aspects of medusae traits were present in the medusozoan common ancestor and which evolved in parallel require the deconstruction of the medusa’s component parts as discussed above. In addition, an understanding of the cellular processes involved in medusa formation will help fill a gap in knowledge between the developmental trajectory and morphology. To this end, we investigated the cell dynamics during medusa development in distinct lineages. Looking at the pattern of cell proliferation, cell migration and programmed cell death, we show that the development of the medusa in representatives of the two main medusozoan lineages relies on distinct cellular behaviors.

## RESULTS

### Study Organisms

In this study we surveyed two scyphozoan species: *Aurelia coerulea* (Discomedusa, Ulmaridae) and *Sanderia malayensis* (Discomedusa, Pelagiidae), and six hydrozoan species: *Craspedacusta sowerbii* (Trachylina, Limnomedusae), *Staurocladia* sp. (Hydroidolina, Cladonematidae), *Hydractinia symbiolongicarpus* (Hydroidolina, Hydractiniidae), *Bouillonactinia carcinicola* (Hydroidolina, Hydractiniidae), *Podocoryna carnea* (Hydroidolina, Hydractiniidae) and *Podocoryna exigua* (Hydroidolina, Hydractiniidae) (Figure 1). These organisms were chosen to represent the diversity of medusae development found across Medusozoa.

**Figure 1:**
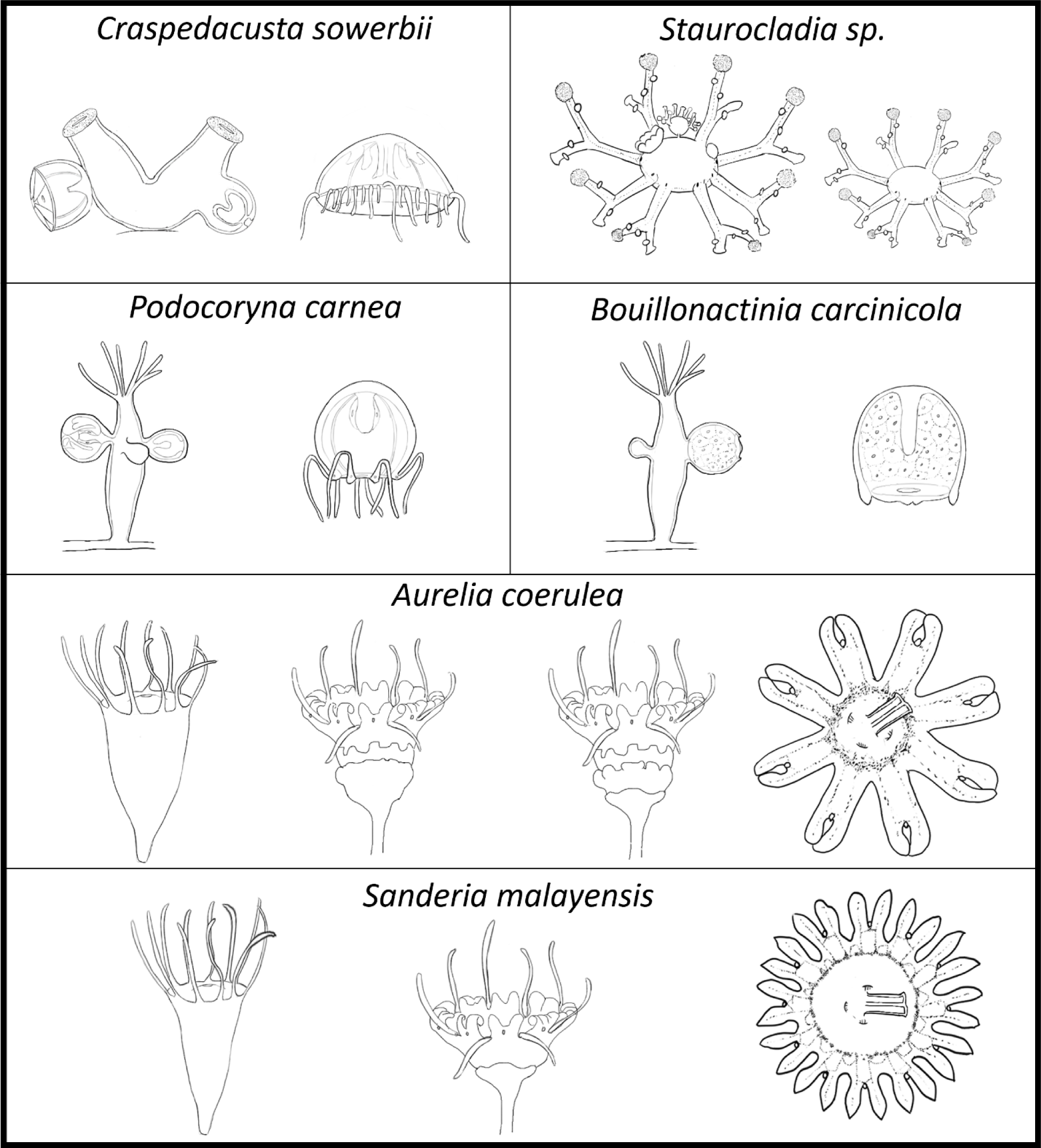
Schematic of the distinct modes of asexual reproduction of the medusa in the study organisms.

Scyphozoans undergo medusa development through a process called strobilation. The first sign of strobilation consists of the formation of a groove below the whorl of tentacles, followed by regression of the polyp tentacles. In polydisk strobilation, such as seen in *Aurelia coerulea* (Figure 1), the oral-most developing disk is ontogenetically more advanced while the aboral most disk exhibits earlier developmental stages. Each disk develops into a juvenile medusa called ephyra. *Sanderia malayensi*s (Figure 1) is a monodisk strobilator. Similar to *A.coerulea* a the first signs of strobilation consist of the formation of a groove below the tentacle whorl and the regression of the tentacles, however only one ephyra is formed.

Unlike scyphomedusae, hydromedusae develop through lateral budding from the polyp and do not transform polyp structures. The first sign of budding consists of the formation of a marginal outgrowth of epithelial cells, followed by the development of the medusa structure in the distal portion of the bud.

The freshwater jellyfish *Craspedacusta sowerbii* is a limnomedusae that belongs to the subclass Trachylina. *C. sowerbii* laterally buds the medusa from either solitary polyps or polyps organized into pseudo-colonies (Figure 1). Each polyp can only produce one medusa and undergoes complete regression after release of the medusa. The first signs of the medusa budding in *C.sowerbii* consist of lateral outgrowth. Early stages of medusa are accompanied by a slight bending of the neck of the polyp head, positioning the medusa bud upright, subsequent regression of the head and ultimately regression of the polyp into a rudiment.

*Podocoryna carnea, Podocoryna exigua*, *Bouillonactinia carcinicola* and *Hydractinia symbiolongicarpus* are all colonial hydrozoans belonging to the family Hydractiniidae. All three were included because they possess medusae and different stages of medusae reduction (eumedusoid and sporosac), respectively, enabling us to address the role in cellular dynamics in the evolutionary reduction of the hydromedusa from closely related lineages.

Medusa development in *Podocoryna carnea* is initiated through lateral budding in a subset of polyps in the colony. Lateral budding takes place in a region of the polyp called the budding zone. *P. carnea* is able to produce several medusa buds, successively, within the budding zone (Figure1). During medusa budding, the budding polyp regresses starting with resorption of the tentacles, closure of the mouth opening and reduction of the hypostome. *Podocoryna exigua* exhibits similar colony architecture and medusa budding development as *P. carnea*. *P. exigua* was used in this study as it spontaneously releases budding and non-budding polyps from the colony enabling manipulation of budding polyps with minimal disruption. By contrast, removal of budding polyps from *P. carnea* colonies and/or injury of the developing medusa leads to resorption of the medusa bud. Unlike *Podocoryna*, *Bouillonactinia carcinicola* does not bud fully developed medusae and instead buds reduced medusae called eumedusoids. The eumedusoid of *B. carcinicola* is short-lived (few hours) and lacks a mouth, true gonads, and tentacles (Figure1). Similar to *Podocoryna*, the eumedusoid budding occurs in a subset of the polyps of the colony and several eumedusoids are budded simultaneously. The eumedusoid budding is accompanied by regression of the oral structures of the budding polyp. *Hydractinia symbiolongicarpus* lacks a medusa and instead buds sporosacs laterally from specialized non-feeding polyps called gonozooids. The sporosac lacks all medusa features and simply consists of a bilayered structure containing the gametes.

*Staurocladia* sp. is a capitate colonial hydrozoan that exhibits the rare ability to asexually bud medusae from the medusa, in addition to laterally budding from polyps. In this study, we did not have access to the polyp stage of *Staurocladia* sp. and instead characterized medusae budding from medusae. The medusa of *Staurocladia*, sometimes referred to as the crawling jellyfish, is a free living, non-swimming medusa with a flattened bell. Although non-swimming, the medusa of *Staurocladia* exhibits all features of a fully developed medusa and was used in this study to address the role of cellular dynamics in the medusa-medusa budding, in contrast to the more common form of polyp to medusa budding. The medusa is able to bud several medusae simultaneously (Figure 1). These buds are located on the exumbrella in the bell margin area, more specifically in the inter-tentacular space.

### Medusa development utilizes broad cell proliferation during early ontogeny and a structure-specific cell proliferation in late ontogenesis

In *A. coerulea*, cell proliferation was initially detected broadly throughout the strobila except in the regressing tentacles (Figure 2A). In *A. coerulea*, the early development of the disk is associated with cell proliferation in the developing lappets and rhopalia in an outer ring-like pattern (Figure 2B). As the disks develop, cell proliferation extends to the whole disk and later shows a higher activity in the gastrodermis and rhopalia (Figure 2B and 2D). After release of the ephyra, cell proliferation is concentrated in the gastric filaments (Figure 2C), and the rhopalia (Figure 2D). In *S. malayensis*, a high density of cell proliferation was observed in the mouth of the developing ephyra of the transforming polyp but not in the regressing tentacles (Figure 2E). In later stages of the developing ephyra dense and broad cell proliferation is observed in the gastrodermis of the ephyra in addition to the regenerating oral end of the polyp (Figure 2F). After release of the ephyra an intense cell proliferation is observed in the rhopalia (Figure 2G and 2H) and the gastrodermis of the manubrium (Figure 2G).

**Figure 2:**
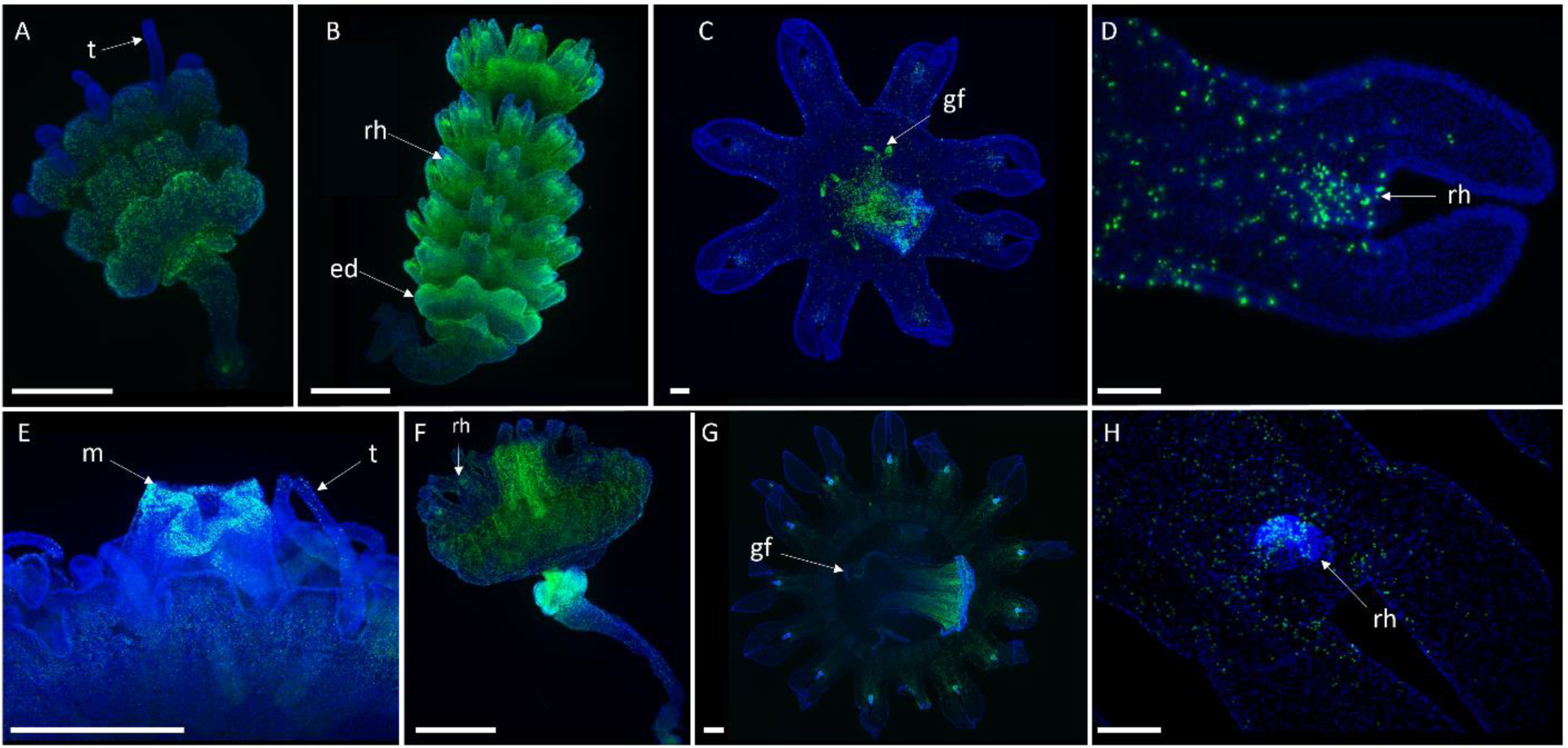
Edu cell labeling in *Aurelia coerulea* (A-D) and *Sanderia malayensis* (E-H), green (Edu) and blue (Hoechst). Early strobila (A), Late strobila (B), newly released ephyra (C), ephyra lappet (D). Developing ephyra mouth (E), late strobila (F), newly released ephyra (G), ephyra lappet (H). ed, early disk; gf, gastric filaments; m, mouth; rh, rhopalium; t, tentacle; Scale bars: 500 μm (A, B, E, F), 100 μm (C, D, G, H).

In the freshwater hydrozoan jellyfish *C. sowerbii*, the initiation of the medusa bud is relatively unpredictable and difficult to detect, thus only the later stages of early budding could be characterized. Cell proliferation is detected at low levels throughout the early medusa and at higher densities in the distal portion of the bud (Figure 3A). In later stages, dense cell proliferation is detected in the developing manubrium and the bell margin (Figure 3B and 3C), however Edu labeled cells are also found in the radial canals (Figure 3C). The final stage of medusa budding shows the highest overall activity of cell proliferation occurring in the manubrium (Figure 3B and 3C), bell margin and developing tentacles (Figure 3C). In the newly released medusa, a high density of cell proliferation is maintained in the manubrium and to a lesser degree in the bell margin (Figure 3D). Edu labeled cells were no longer observed in the radial canals but were detected in the bell margin area (Figure 3D).

**Figure 3:**
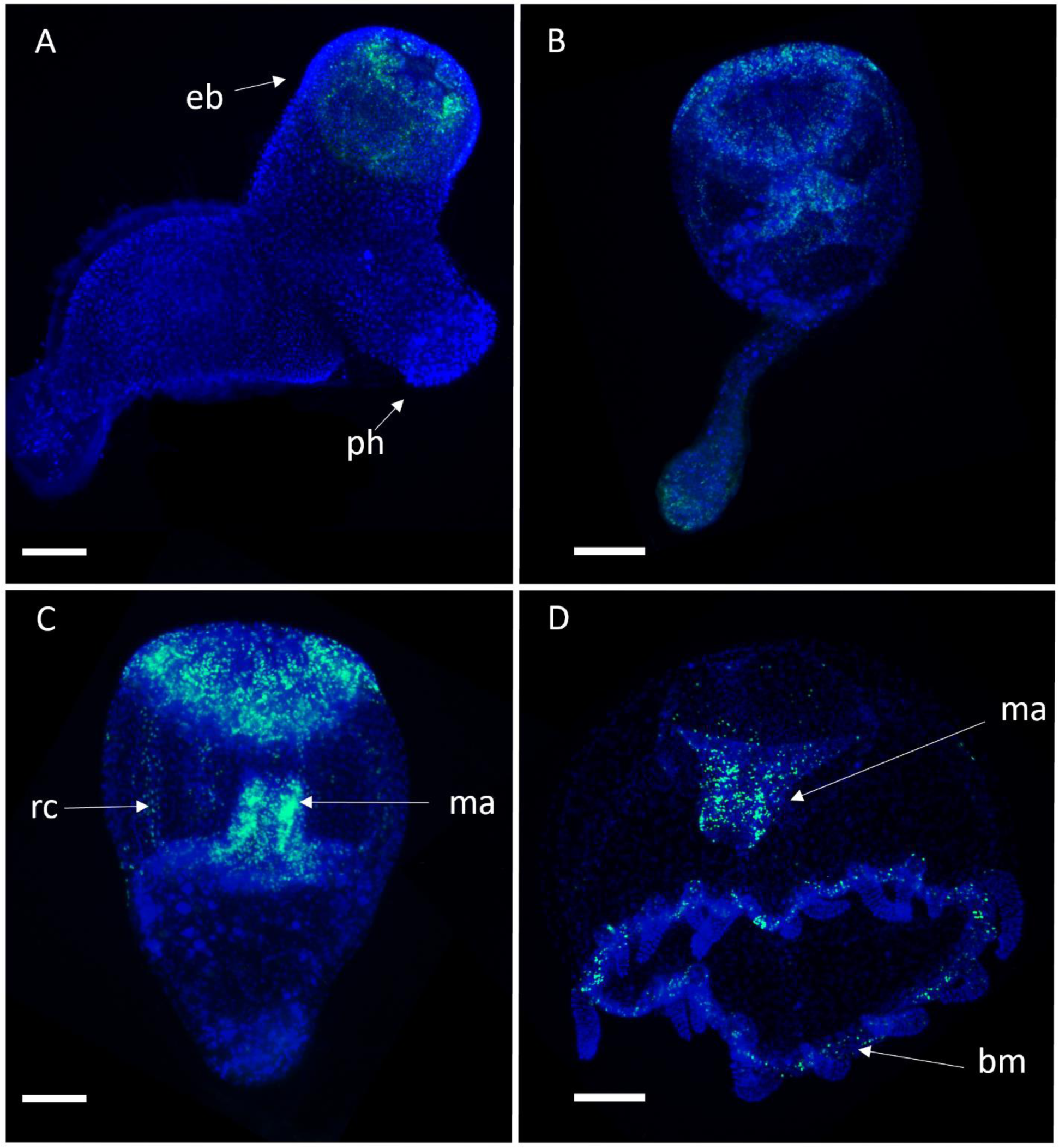
Edu cell labeling in *Craspedacusta sowerbii* (A-D), green (Edu) and blue (Hoechst). Early bud (A), intermediate bud (B), late bud (C), newly released medusa (D). eb, early bud; ma, manubrium; ph, polyp head; rc, radial canal. Scale bars: 200 μm (A-D)

Three species of hydractiniid hydrozoans that have different types of gonophores were examined; *H. symbiolongicarpus* lacks a medusa and instead has a sporosac; *P. carnea* has a medusa state in its life cycle; *B. carnicola* has a reduced medusa, or medusoid. In *H. symbiolongicarpus*, Edu labeled cells were found scattered in the epidermis of the sporosacs (Figure 4A). In the early buds of *P*. *carnea*, dense and broad cell proliferation was detected (Figure 4B). In intermediate medusa buds, a dense population of proliferating cells was detected, concentrated in the distal portion of the bud (Figure 4B). In the latest stages of medusa bud development, cell proliferation was restricted to the distal-most portion of the bud where the various medusa structures are developing (e.g. the velum, tentacles, ring canal, nerve rings) (Figure 4B). After the release of the medusa, cell proliferation was detected exclusively in the manubrium, the tentacle bulbs, and the proximal portion of the tentacles (Figure 4C). Early medusa buds of *B. carcinicola* showed intense and broad cell proliferation (Figure 4D, 3E) while later stages showed an increasingly distally restricted cell proliferation pattern towards the distal end of the bud (Figure 4D, 4E). The released medusoid of *B. carcinicola* did not exhibit any cell proliferation in the rudimentary tentacle bulbs, nor in the manubrium (Figure 4F).

**Figure 4:**
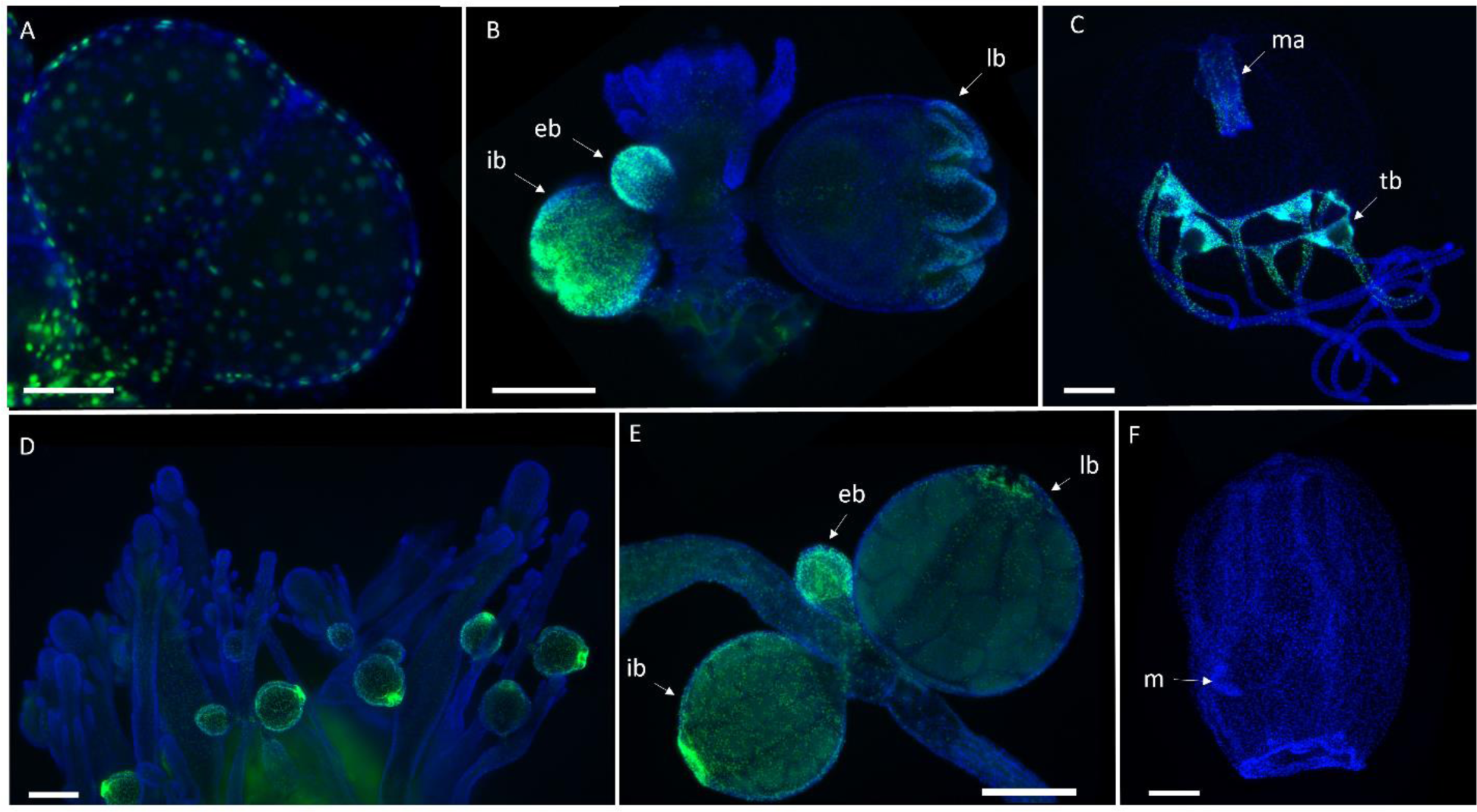
Edu cell labeling of three hydractiniid species, *Hydractinia symbiolongicarpus* (A), *Podocoryna carnea* (B-D), *Bouillonactinia carcinicola* (E-G), green (Edu) and blue (Hoechst). Female sporosacs (A), feeding polyp (B), budding polyp (C), newly released medusa (D), colony exhibiting feeding and budding polyps (E), budding stages (F), newly released medusoid (G). eb, early bud; ib, intermediate bud; lb, late bud; m, mouth; ma, manubrium; tb, tentacle bulb. Scale bars: 200 μm (A-F).

The medusa of *Staurocladia* sp. is capable of asexually budding medusae from its outer bell. In the budding medusa of *Staurocladia* sp., Edu labeled cells were detected at the base of the bud in the bell (central disk) margin, and in the early medusa bud (Figure 5A). In intermediate stages of the medusa bud, cell proliferation was detected in the developing tentacles (Figure 5B). After release of the medusa, Edu labeled cells were found scattered across the newly released medusa and more specifically in developing tentacles (Figure 5C).

**Figure 5:**
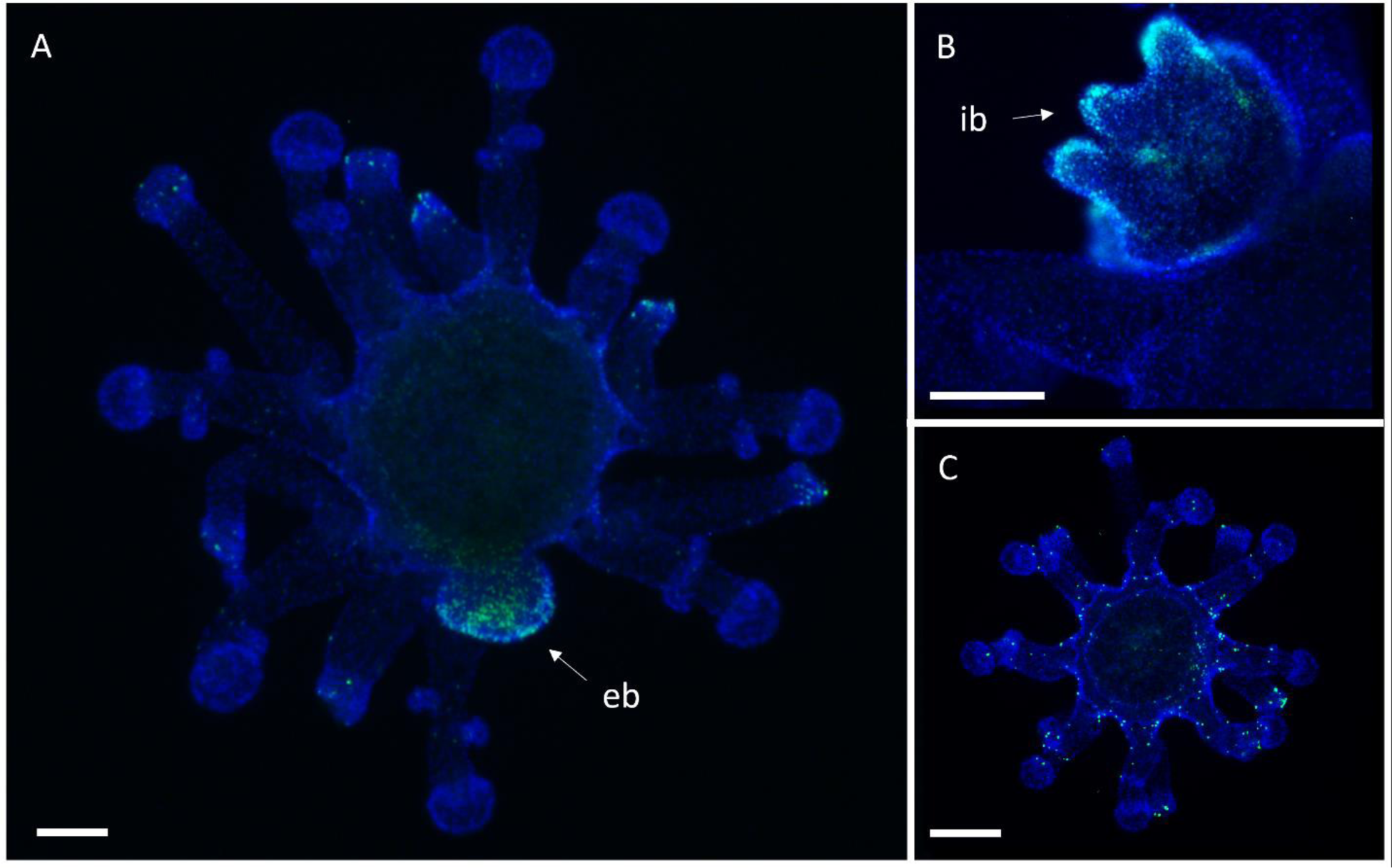
Edu cell labeling of *Staurocladia* sp. (A-C), green (Edu) and blue (Hoechst). Medusa bearing an early bud (A), intermediate medusa bud (B), newly released medusa (C). eb, early bud; ib, intermediate bud. Scale bars: 200 μm (A-C).

### Cell tracing reveals differences in cell recruitment dynamics between scyphomedusa and hydromedusa development

Dil, a fluorescent lipophilic dye, was used to label cells to trace their movement during medusa production. A patch of cells was labeled below the developing disks in *A. coerula* (Figure 6A) and *S. malayensis* (Figure 6F). Neither show a pattern of migration throughout strobilation. Labeled cells are found within the same area of the body column of the polyp 24h after injection (n=18/18) (Figure 6B and 6G). In *A. coerulea*, the position of the labeled cells corresponded to newly formed disks (n=18/18) (Figure 6B). 48 hours after cell labeling, no DiI positive cells were observed in the strobila of *A. coerulea* (n=18/18) (Figure 6C). In *S. malayensis*, labeled cells remained at the site of the injection in most animals (n=12/18) (Figure 6H). In the remaining animals (n=6/18) no labeled cells were detected (not shown). After the release of the ephyra of *S. malayensis*, labeled cells were still found in the body column of the regenerated polyp (n=10/18) (Figure 6I) in the remaining animals (n=8/18) no labeled cells were detected.

**Figure 6:**
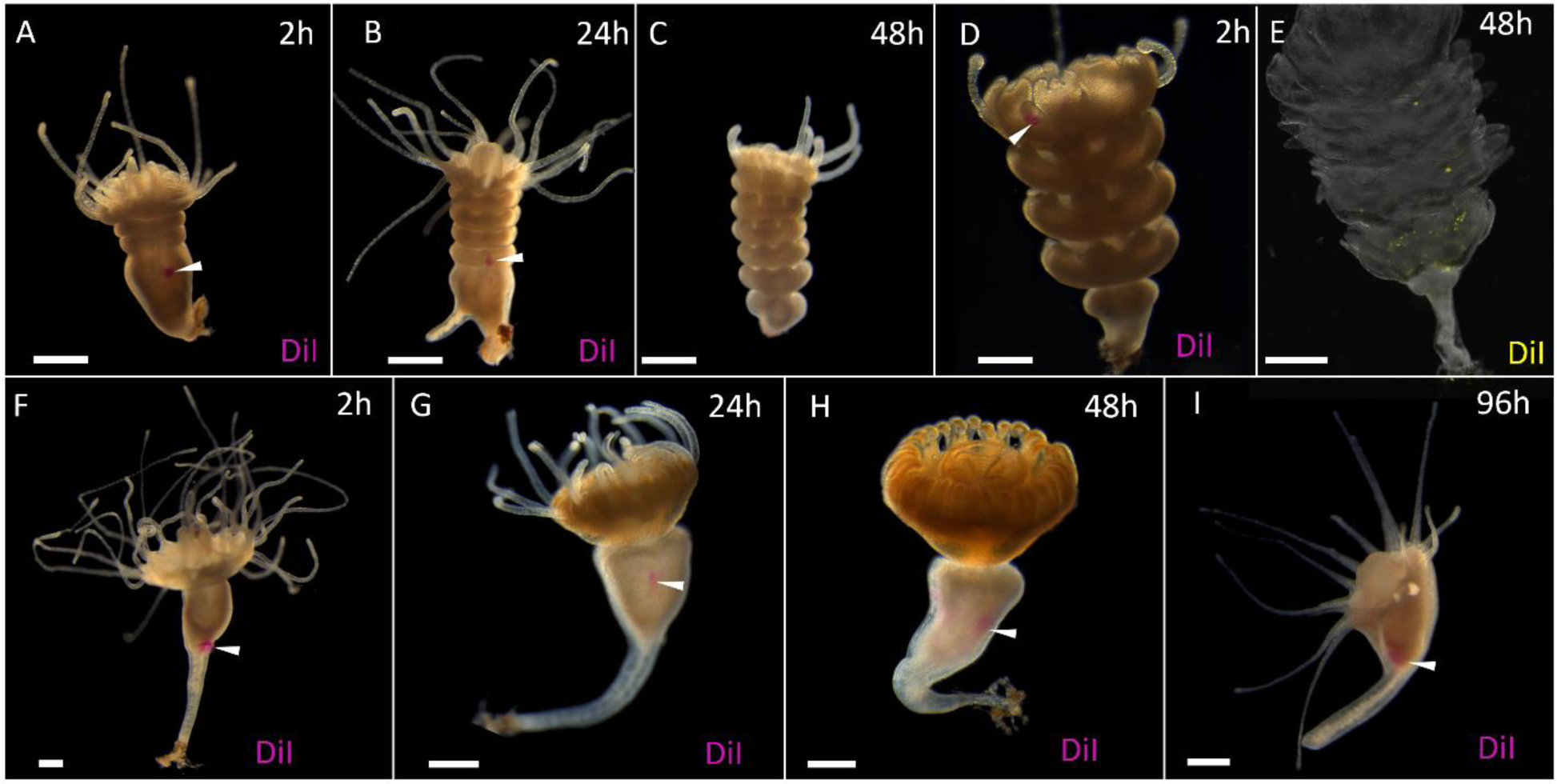
DiI cell tracing during strobilation in two scyphozoan species, *Aurelia coerulea* (A-E) and *Sanderia malayensis* (F-I). Early strobila 2h after body column cell labeling via DiI injection (A), Early strobila 24h after body column cell labeling via DiI injection (B), late strobila 48h after body column cell labeling via DiI injection (C). Late strobila 2h after regressing tentacles cell labeling via DiI injection (D), Late strobila 48h after regressing tentacles cell labeling via DiI injection (E). Early strobila 2h after body column cell labeling via DiI injection (F), Early strobila 24h after body column cell labeling via DiI injection (G), late strobila 48h after body column cell labeling via DiI injection (H), regenerated polyp after release of the ephyra 96h after body column cell labeling via DiI injection (I). Scale bars:1mm.

During strobilation, radial and ad-radial tentacles regress. Radial tentacles regress and form the rhopalia in the oral most developing ephyra (Berrill, 1949), while the contribution of the ad-radial tentacles is unknown. To assess the fate of the ad-radial tentacles, regressing ad-radial tentacles of *A. coerulea* were labeled with Dil (Figure 6D). 48 hours after cell labeling, labeled cells appeared to migrate from the most apical developing ephyra to the most basal, and sporadically in intermediate developing ephyrae (n=12/12) (Figure 6E). In *S. malayensis*, the same experiment was carried out, however injected tentacles were either shed by the animals or no signal was detected 24h after injection (not shown).

To assess the contribution of the polyp tissue to the development of the hydromedusa, cells from the body column of the budding hydroids of *Podocoryna exigua* and *C. sowerbii* were labeled by injection of DiI. *P. exigua* was chosen as an alternative to *P. carnea*, as removal from the colony and injection caused regression of the medusa bud in *P. carnea*. In contrast, *P. exigua* spontaneously releases budding polyps from the colony and the development of the medusa showed negligible impairment from injection of DiI.

In *C. sowerbii*, cells were labeled below the budding zone bearing early medusa buds (Figure 7A). After 48h, labeled cells were observed in the intermediate medusa bud, more specifically in the developing manubrium, radial canals, the ring canal with a few scattered labeled cells in the epidermis of the bell (n=12/12) (Figure 7B). In the newly released medusa, labeled cells were still found in the manubrium, the radial canals, and the ring canal (n=8/12) (Figure 7C).

**Figure 7:**
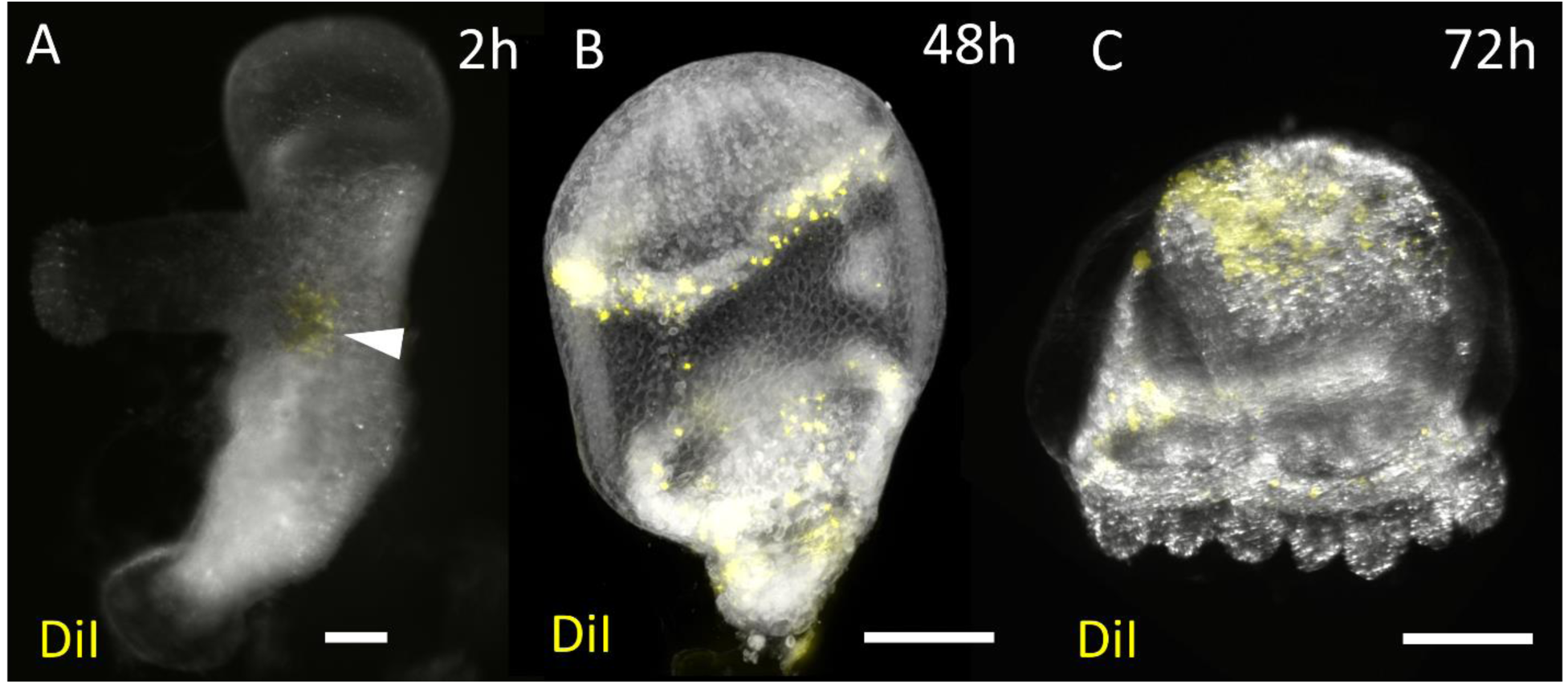
DiI cell tracing medusa budding in *Craspedacusta sowerbii* (A-C). Early budding polyp 2h after body column cell labeling via DiI injection (A), intermediate bud 48h after body column cell labeling via DiI injection (B) and newly released medusa 72h after body column cell labeling via DiI injection (C). scale bars: 100μm (A), 200μm (B and C). The white arrowhead indicates the site of injection.

In *P. exigua*, cells were labeled below the budding zone bearing various stages of medusa buds (Figure 6A). After 48h, labeled cells were still observed in the body column of the polyp, with some cells in a more aboral position (n=12/12) (Figure 8B). In addition, labeled cells were found in the tentacles of the budding polyp (n=12/12), and a few cells were found in the developing medusa buds (n=7/12) (Figure 8B). After three days, a dense population of labeled cells were observed in a crescent pattern below the budding region with a higher density being detected at the site of the injection (Figure 8C). In the suboral area of the polyp, previously bearing a whorl of tentacles (Figure 8C), labeled cells were observed. Lastly, a large population of labeled cells was found in the medusa buds and the peduncle connecting the polyp to the medusa (n=12/12) (Figure 8C).

**Figure 8:**
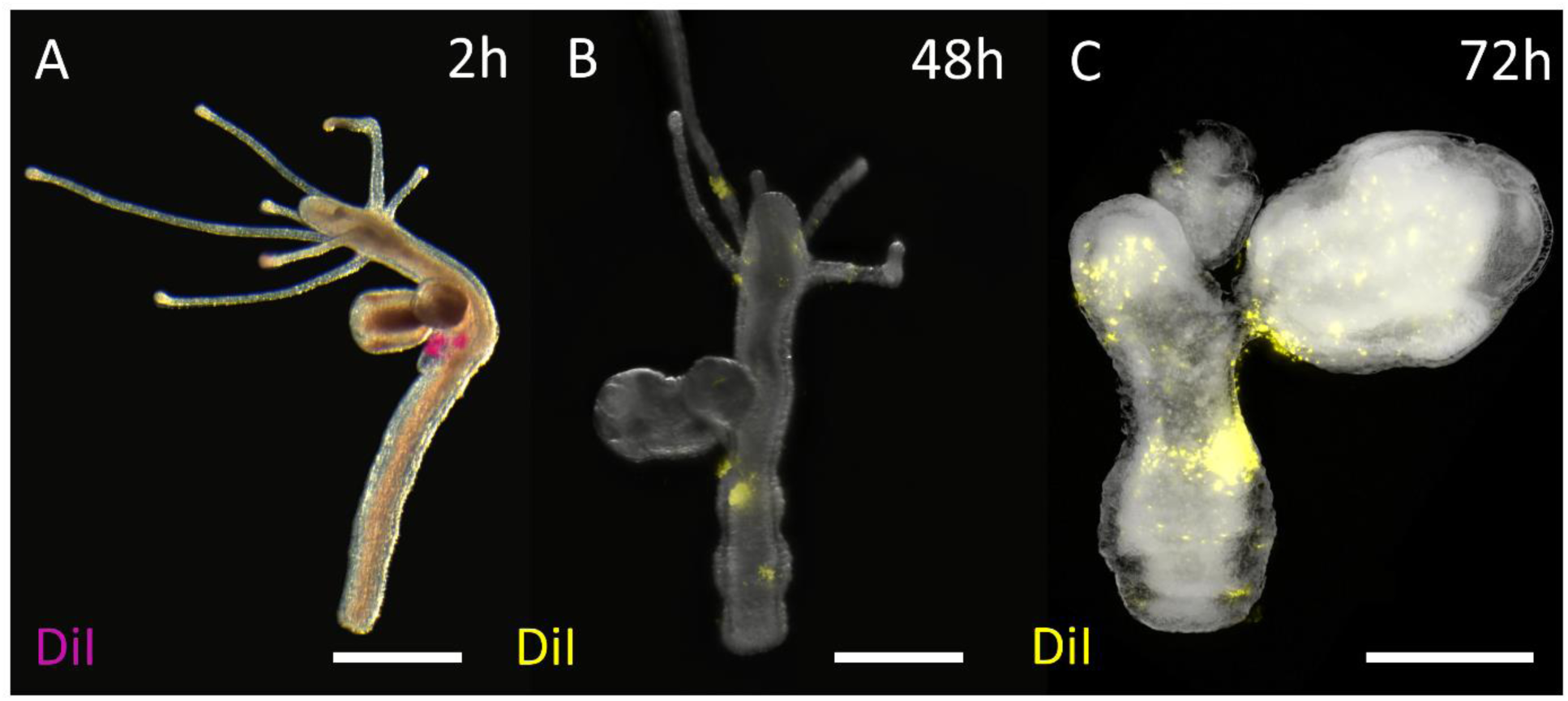
DiI cell tracing medusa budding in *Podocoryna exigua* (A-C). budding polyp 2h after body column cell labeling via DiI injection (A), budding polyp 48h after body column cell labeling via DiI injection (B) late stages of budding 72h after body column cell labeling via DiI injection (C). scale bars: 200μm.

In the budding medusae of *Staurocladia sp*., cells were initially labeled in the central disk of the medusa (Figure 9A). After 48h, labeled cells were observed scattered in the central disk and in the developing medusa bud, more specifically in the umbrella of the medusa bud (n=24/24) (Figure 7B). After three days, labeled cells were found almost exclusively in the medusa bud, in the umbrella and additionally in the developing tentacles (n=24/24) (Figure 9C).

**Figure 9:**
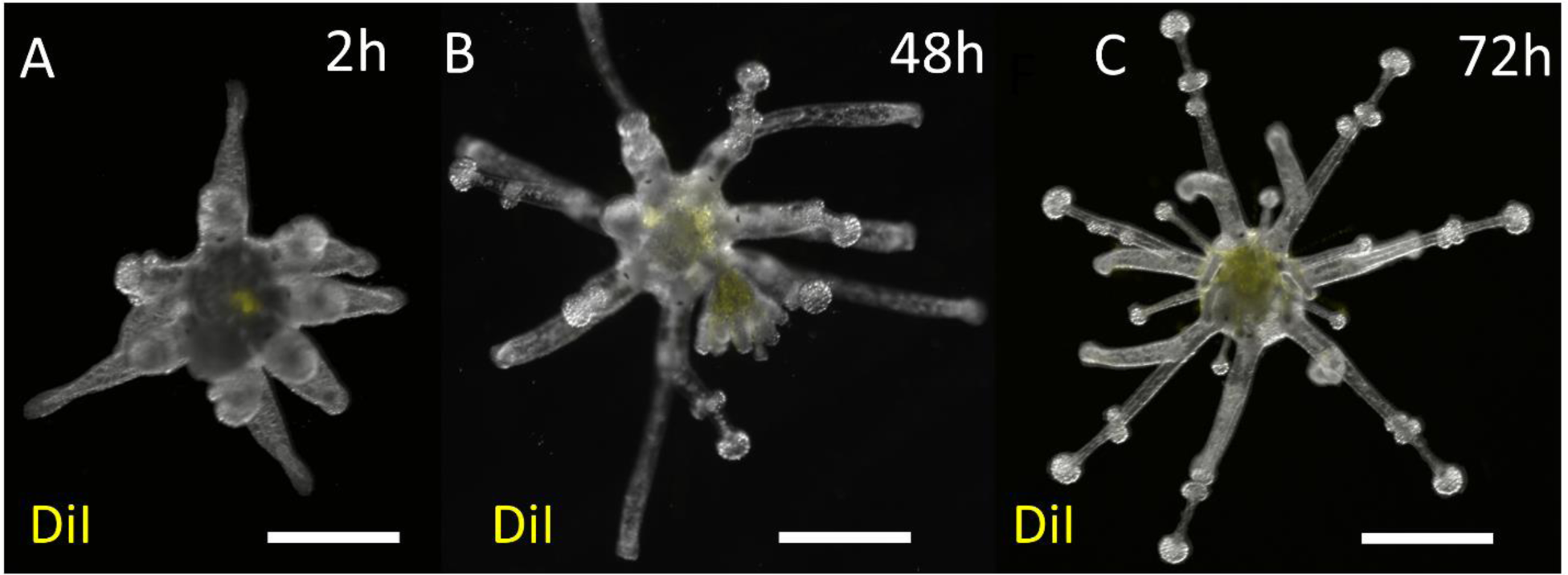
DiI cell tracing medusa budding in *Staurocladia* sp. (A-C). Early budding medusa 2h after exumbrella cell labeling via DiI injection (A), intermediate bud 48h after exumbrella cell labeling via DiI injection (B) late stage of medusa bud (oral view) 72h after exumbrella cell labeling via DiI injection (C). scale bars: 500μm.

### Scyphomedusa development relies on apoptosis

In *A. coerulea*, a dense population of TUNEL positive cells was detected in the early strobila (Figure 10A). It appears that apoptosis is most prevalent in the oral most developing ephyra, more specifically in the developing bell and the tentacles, while a lesser signal is detected in the disks below (Figure 10A). In *S. malayensis*, the TUNEL positive cells were also detected in the early strobila but appeared to be restricted to the tip of the regressing tentacles (Figure 11A), where a lesser signal was detected in later stages of the strobila (Figure 11B, 11C and 11D). In contrast, late stage strobilae of *A. coerulea* exhibited a relatively dense population of TUNEL positive cells throughout the strobila (Figure 10B and 10C). Despite the TUNEL positive cell population density differing between the two species during strobilation, apoptotic cells were detected throughout the epidermal tissue of the developing medusa for both species (see below). The foot of the scyphistoma exhibits autofluorescence, thus the signal did not reflect apoptosis due to the absence of apoptotic bodies in that location.

**Figure 10:**
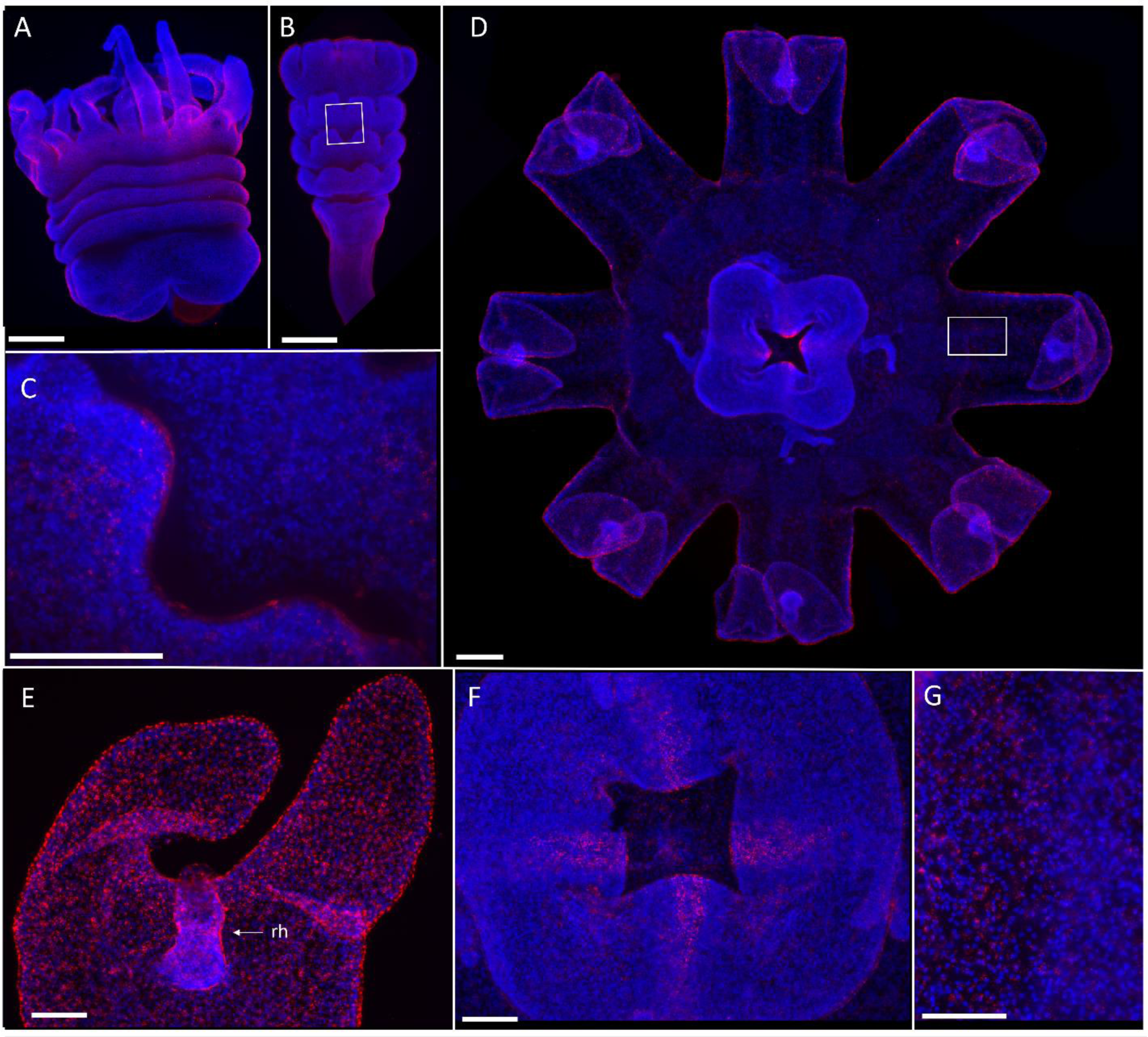
TUNEL assay in *Aurelia coerulea* (A-D), red (TUNEL), blue (Hoechst). Early strobila (A), Late strobila (B), higher magnification of a late strobila(C), newly released ephyra (D), lappet of the ephyra (E), mouth of the ephyra (F), higher magnification of the coronal region of the ephyra(G). Scale bars: 200 μm (A, B, D), 100 μm (C, E, F, G).

**Figure 11:**
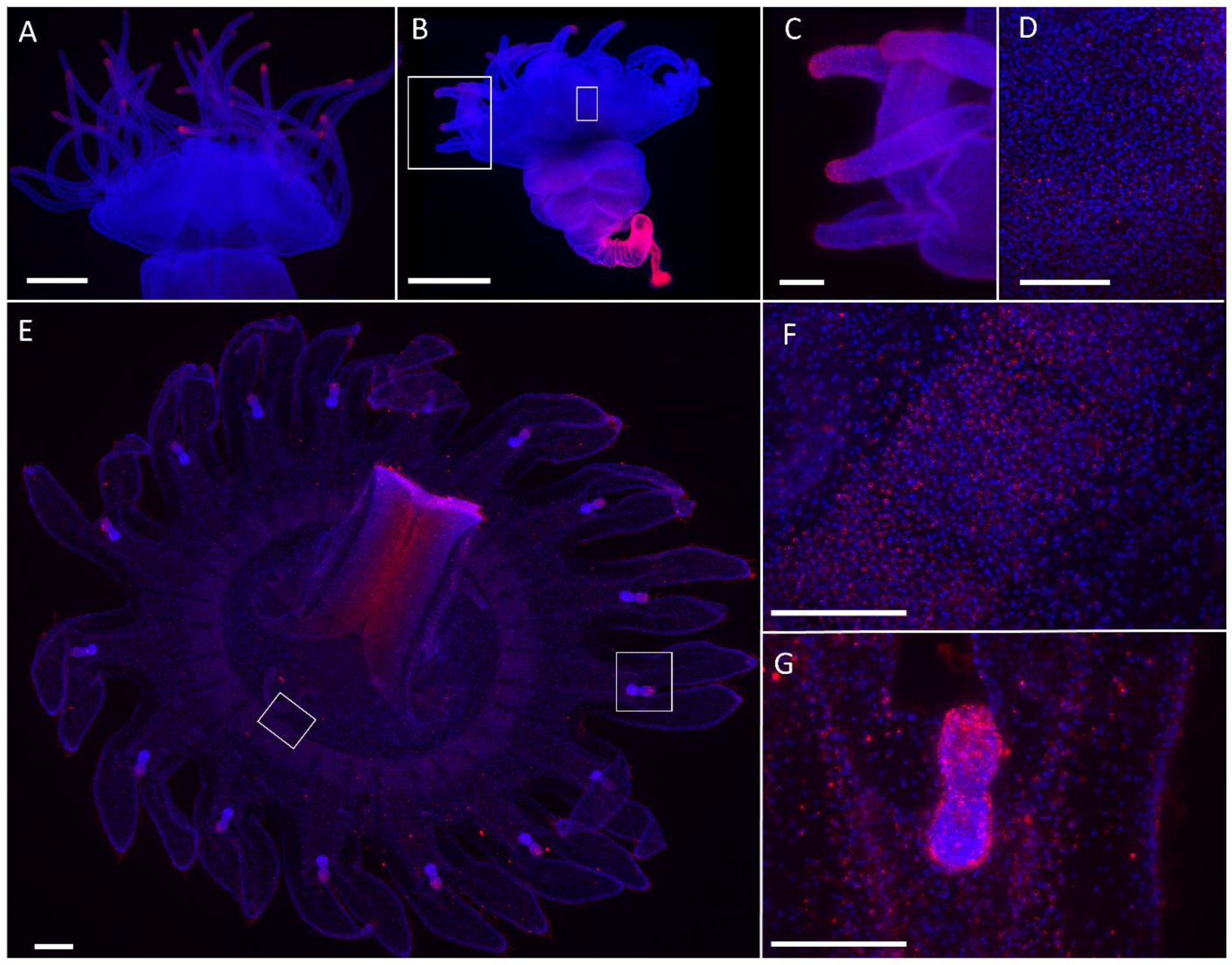
TUNEL assay in *Sanderia malayensis* (A-D), red (TUNEL), blue (Hoechst). Polyp head in a strobila, 96h after indole induction (A), strobila 120h after indole induction (B), higher magnification of (B), in the regressing tentacles of the strobila(C), higher magnification of the regressing tentacles shown in (B),higher magnification of the developing bell of the ephyra shown in(B) (D), newly released ephyra (E), higher magnification of (E), in the coronal region of the ephyra(F), higher magnification of (E), in the lappet region(G). Scale bars: 200 μm (A-B), 100 μm (D-G).

In the released ephyra, an apoptosis signal was detected throughout (Figure 10D, 10G, 11E and 11F), with a dense signal in the manubrium and the rhopalia (Figure 10D, 10E, 10F, 11E, 11F and 11G). Although apoptotic cells were almost exclusively restricted to the epidermis in the strobilae in *A. coerulea*, the released ephyra exhibited apoptotic cells in both gastrodermis and epidermis. A distinct radial apoptosis pattern was observed in the manubrium of the ephyra (Figure 10F) and extended down the gastrodermis of the manubrium (Figure 11E). Additionally, apoptotic cells were detected in the coronal area of the ephyra (Figure 10G and 11F).

In the five hydrozoan species tested (*H. symbiolongicarpus*, *B. carcinicola*, *P. carnea*, *C. sowerbii* and *Staurocladia* sp.), no TUNEL positive cells were detected during gonophore development, with one exception. After spawning, the medusoid of *B. carcinicola* exhibited an intense apoptosis signal throughout the medusoid (Figure S1D and S1E).

### Effect of inhibiting cell division and programmed cell death on the development of the medusa

Hydroxyurea is an antimetabolite that inhibits cell division by preventing cells from leaving the G1/S phase of the cell cycle. Z-VAD-FMK is a cell-permeable pan-caspase inhibitor that irreversibly binds to the catalytic site of caspase protease and thus can inhibit initiation of apoptosis. To address the function of apoptosis and cell proliferation during medusa development, two scyphozoan representatives (*A. coerulea*, *S. malayensis*) and three hydrozoan representatives (*P. carnea*, *C. sowerbii* and *Staurocladia* sp.) were treated with hydroxyurea and Z-VAD-FMK, separately during medusa development.

After chemical induction of strobilation using indole (Helm and Dunn, 2017), the first signs of strobilation in *A. coerulea* were observed after 48h. Animals treated with 10µM hydroxyurea showed effective inhibition of cell division (Figure 12B). Most of the animals (86%) treated with hydroxyurea were able to initiate strobilation, while all the polyps in the control group underwent strobilation (Figure 12E). Strobilae from treated animals produced a reduced number of disks per polyp (1.23 ± 0.13, p<0.0001) compared to the control group (4.38 ± 0.28) (Figure 12F), which did not develop into ephyrae (Figure 12G). After five days of treatment, strobilae underwent reverse development, the disks resorbed, and the polyp tentacles ultimately regenerated (Figure 13). This reversal is associated with a reduction of the number of disks (0.38 ± 0.15) (Figure 12F and 13). In *A. coerulea* treated with Z-VAD-FMK, animals showed effective inhibition of apoptosis (Figure 12D) and were not able to initiate strobilation within five days unlike the control group (p<0.001) (Figure 12E, 12F and 13). In treated animals, none of the induced polyps’ tentacles regressed into rhopalia and disks did not form (Figure 12D and 13), thus no ephyra was released (Figure 12G and 13). After a week of treatment, disks were fully resorbed and reversal to polyp was complete (Figure 13). As the use of 10µM hydroxyurea resulted in significant hindering of strobilation and complete inhibition of medusa development, a treatment using 1µM was carried out to assess the impact of cell proliferation in the developing ephyra. In all animals treated with 1µM hydroxyurea, strobilation was initiated and development into ephyrae was successful (Figure 13). The ephyrae, however, exhibited an altered morphology. The lappets were shorter and connected at a wider angle to the lappet stem than the control group (Figure 13). The rhopalia lacked pigment spots and instead exhibited prominent terminal segments bearing large lithocysts (Figure S2). The aboral most disk had undergone reversal of the development and lithocysts in the distal end of the polyp tentacles were observed (Figure 13).

**Figure 12:**
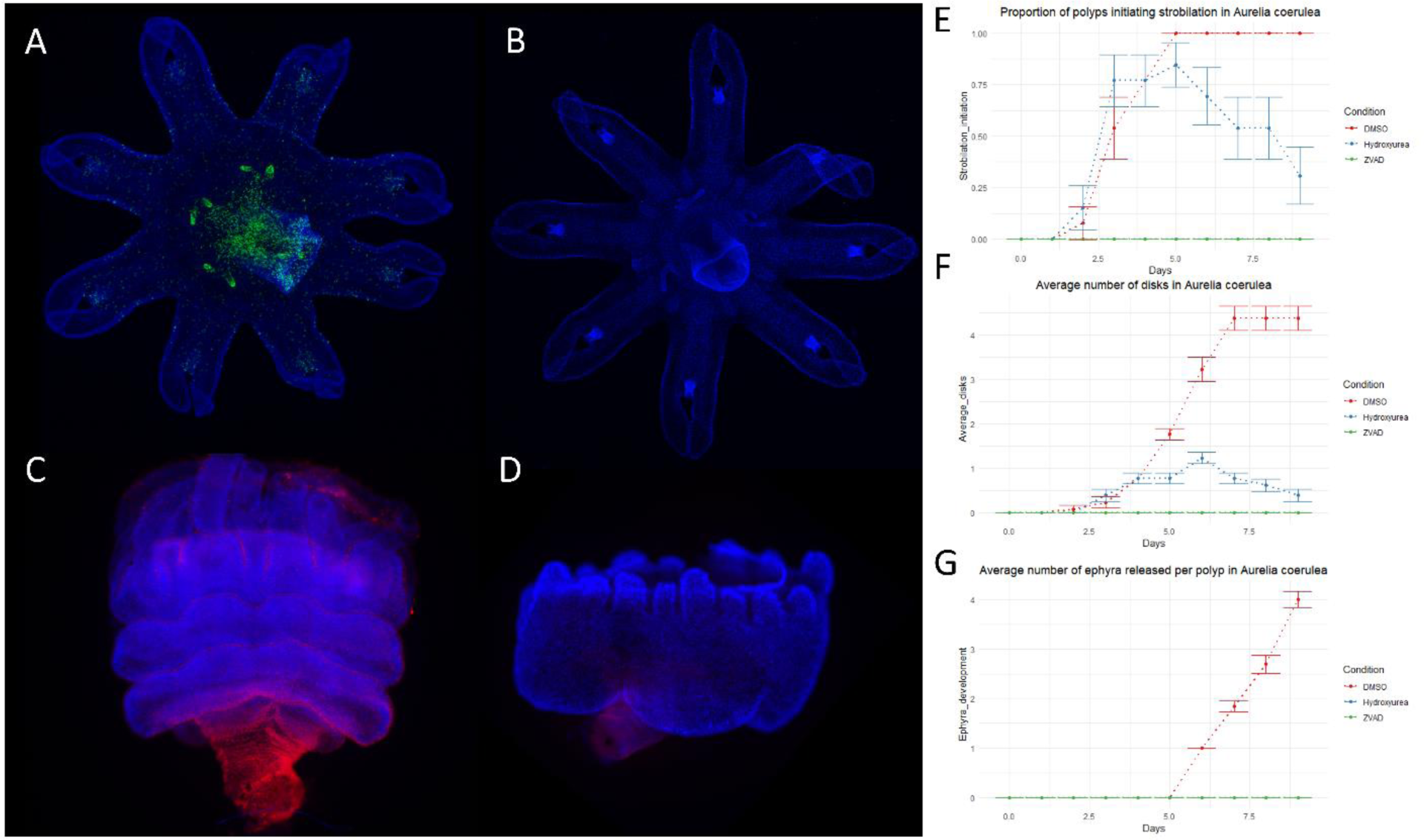
Effect of cell proliferation inhibition (hydroxyurea 10 µM) and cell apoptosis inhibition (Z-VAD-FMK 100 µM) on the development of *Aurelia coerulea*. Edu cell labeling on ephyra incubated in 0.5% DMSO for 48h (A). Edu cell labeling on ephyra incubated in 10µM hydroxyurea for 48h. Green (Edu) and blue (Hoechst) (B). TUNEL-assay, on strobila incubated in 0.5% DMSO for 48h (C), in 100 µM Z-VAD-FMK (D), red (TUNEL), blue (Hoechst). Proportion of polyp initiating strobilation (E). Average number of disks per strobila (F). Average number of ephyrae released per strobila, since no ephyra were released in neither hydroxyurea nor Z-VAD-FMK treated animals, tracing on the graph overlap (G).

**Figure 13:**
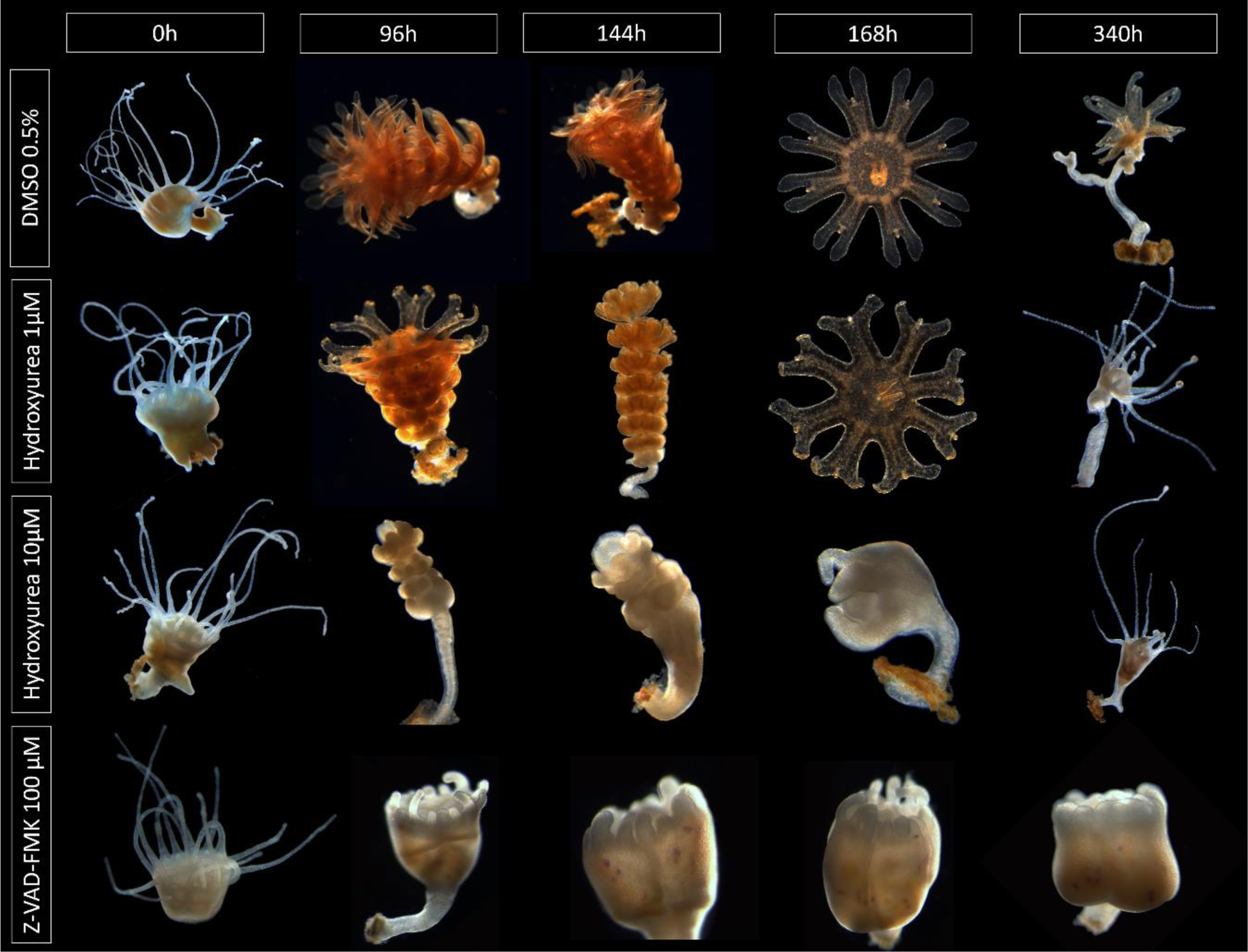
Strobilating individuals of *Aurelia coerulea*, treated with hydroxyurea (1uM and 10 µM) or Z-VAD-FMK 100 µM. Images presented are not in scale.

*S. malayensis* polyps were induced in indole and the first signs of strobilation were observed after 48h. Animals treated with 10µM hydroxyurea showed effective inhibition of cell division (Figure 14B). Most of the animals (72%) treated with hydroxyurea at 0h were unable to initiate strobilation, while all animals in the control group underwent strobilation (Figure 14E). Similar to *A. coerulea*, *S. malayensis* underwent reversal of the medusa development after six days of hydroxyurea treatment when treated prior to the first signs of strobilation (Figure 15). Typically, strobilation in *S. malayensis* starts with the formation of a groove below the polyp head in addition to the regression of the tentacles. As the head transforms and develops into an ephyra, new polyp tentacles develop below the groove. These tentacles belong to the regenerating polyp head, bound to remain after the release of the ephyra. In the treated animals, the groove as well as the second set of tentacles were formed, however, an ephyra was not formed (Figure 15). After a week, the groove resorbed, and the second set of tentacles remained. Similar to the experiment carried out for *A. coerulea*, *S. malayensis* were treated with 1µM hydroxyurea, however, no significant impact on the strobilation was observed (not shown). To address the impact of cell proliferation on the ephyra development, animals were treated with 10µM hydroxyurea at different stages of strobilation instead.

**Figure 14:**
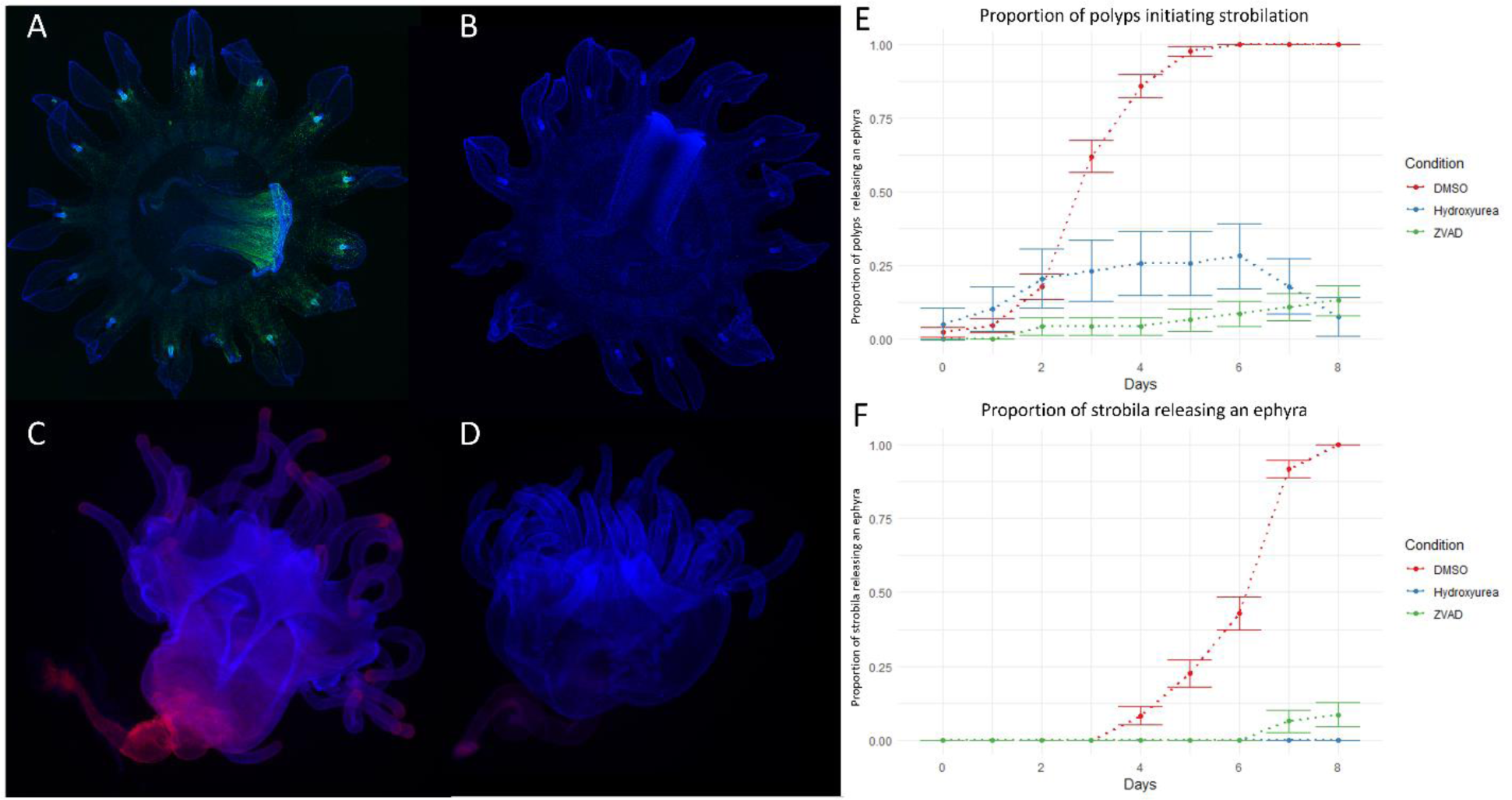
Effect of cell proliferation inhibition (hydroxyurea 10 µM) and cell apoptosis inhibition (Z-VAD-FMK 100 µM) on the development of *Sanderia malayensis*. Edu cell labeling on ephyra incubated in 0.5% DMSO for 48h (A), green (Edu) and blue (Hoechst). Edu cell labeling on ephyra incubated in 10µMhydroxyurea for 48h (B), green (Edu) and blue (Hoechst). TUNEL-assay, on strobila incubated in 0.5% DMSO for 48h (C), in 100 µM Z-VAD-FMK (D), red (TUNEL), blue (Hoechst) (D). Proportion of polyp initiating strobilation (E). Average number of ephyrae released per strobila (F).

**Figure 15:**
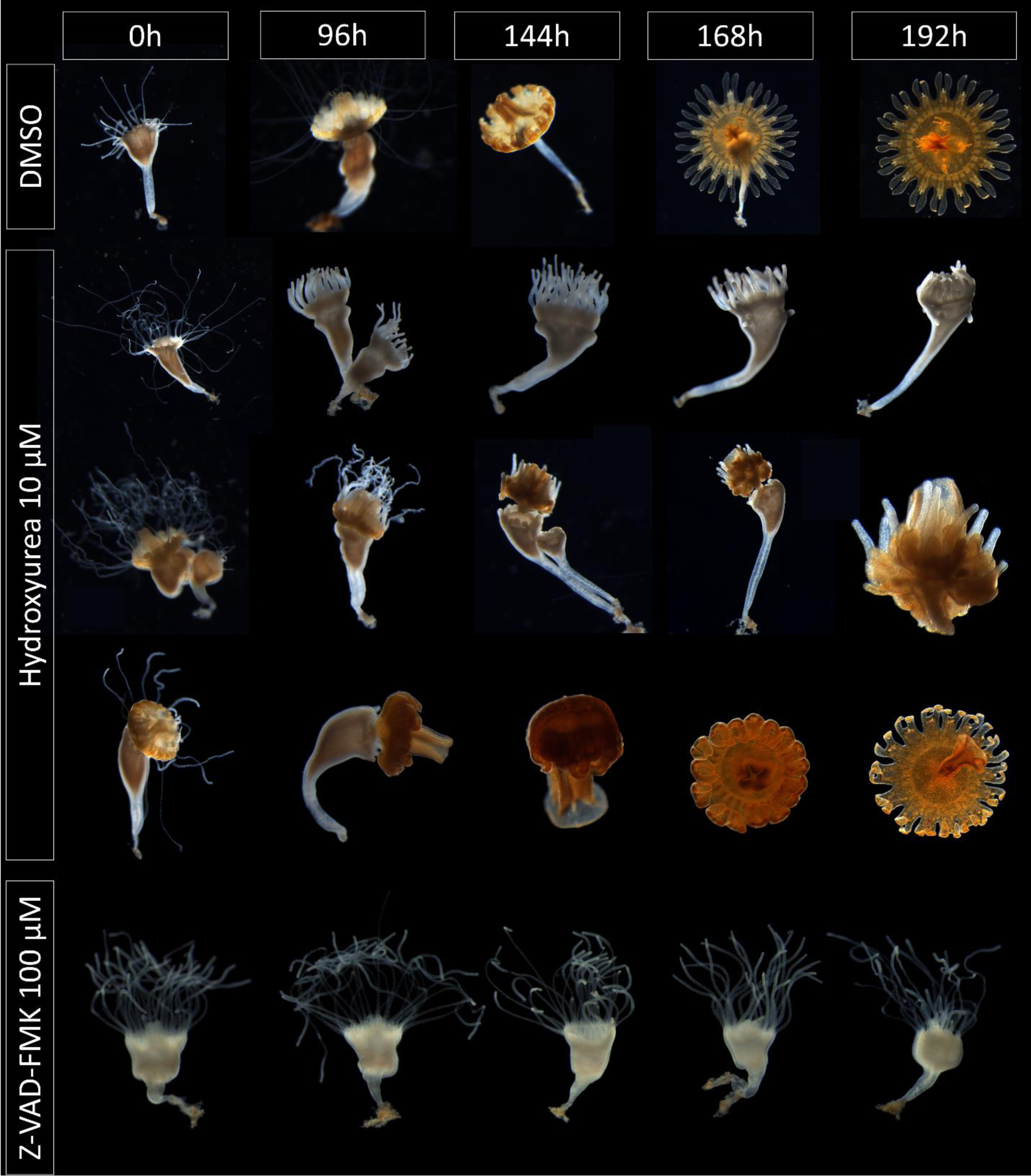
Strobilating individuals of *Sanderia malayensis*, treated with hydroxyurea (10 µM and 1µM) or Z-VAD-FMK 100 µM. Images presented are not in scale. The top row of the 10 µM hydroxyurea treatment corresponds to exposure to hydroxyurea 48h after induction by indole. The middle row of the 10 µM hydroxyurea treatment corresponds to exposure to hydroxyurea 96h after induction by indole. The bottom row of the 10 µM hydroxyurea treatment corresponds to exposure to hydroxyurea 120h after induction by indole.

After four days of induction in indole, animals were treated with 10µM of hydroxyurea. In this experiment, the medusae exhibited a very altered morphology (Figure 15). Complete tentacle regression, development of the bell and medusa structures development did not occur (Figure 15). After 8 days, the underdeveloped medusae were released (Figure 15). After six days of induction in indole, animals were treated with 10µM of hydroxyurea. In this experiment, the ephyrae exhibited a mild alteration of their morphology. Complete tentacle regression and development of medusae structures were achieved. In the released ephyrae, lappets were shorter, and the gastric cavity underdeveloped. *S. malayensis* ephyrae do not naturally possess pigment spots, thus no major morphological alterations of the rhopalia were observed.

In *S. malayensis* treated with Z-VAD-FMK, almost none of the animals (13%, p<0.0001) were able to initiate strobilation throughout the whole experiment (Figure 14E and Figure 15). Consequently, the treated animals produced almost no ephyrae (8.7%, p<0.0001) (Figure 14F).

In *C. sowerbii*, newly released medusae treated with 10 µM hydroxyurea showed effective inhibition of cell division (Figure 16B). *C. sowerbii* polyps only bud one medusa. In all the animals from the control group medusa development was completed within 48h (Figure 16H. At 48h all early and intermediate buds developed into free-living medusae in the control group (Figure 16G, 16H). In animals treated with hydroxyurea, the average number of polyps producing early buds shows a steep reduction over the first 72h, from 0.76 ± 0.10 to none (Figure 16F). During the first 24h, the number of intermediate medusa buds developing from early buds increases and then reduces over a period of three days until none remain, presumably because no buds could be initiatied (Figure 16G). Throughout the experiment only 38% of the buds developed into medusae in the treated animals, which developed from intermediate buds initially present at the start of the experiment (Figure 16H). Released medusae did not exhibit any major alteration of their morphology. In treated animals, 52% of all buds underwent regression (Figure 16I). The slow decrease of intermediate and late bud stages from 48h to 96h corresponds to the regression of the bud (Figure 16G). Increase of early buds at 96h corresponds to the last stage of regression (Figure 16F).

**Figure 16:**
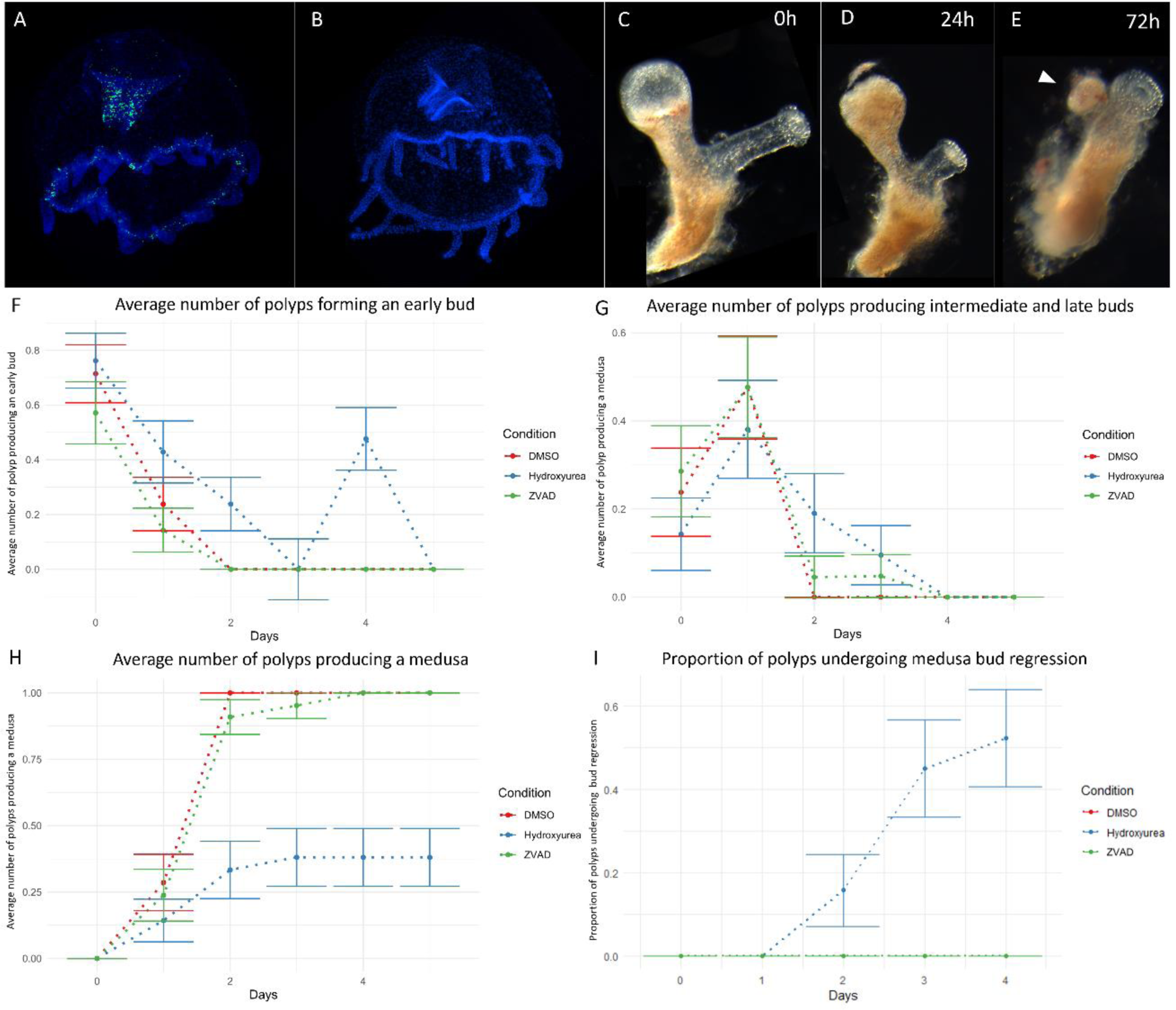
Effect of cell proliferation inhibition (hydroxyurea 10 µM) and cell apoptosis inhibition (Z-VAD-FMK 10 µM) on the development of *Craspedacusta sowerbii*. Edu cell labeling on a newly released medusa incubated in 0.2% DMSO for 48h (A), green (Edu), and blue (Hoechst). Edu cell labeling on a newly released medusa incubated in 10µM hydroxyurea for 48h (B), green (Edu) and blue (Hoechst). Early budding polyp treated with 10 µM hydroxyurea at 0h (C). Budding polyp treated with 10 µM hydroxyurea at 24h, showing regression of the bud (D). Budding polyp treated with 10 µM hydroxyurea at 48h, showing late stages of regression of the bud (E). Average number of polyps producing early buds (F) Average number of polyps producing intermediate and late buds (G). Average number of polyps producing a medusa (H). Percentage of total buds undergoing regression (I). The white arrowhead indicates the last stage of regression of the medusa bud.

*P. carnea* budding polyps can bud several medusae successively, thus an individual polyp bears multiple medusa buds of various stages simultaneously. Animals treated with 10µM hydroxyurea showed effective inhibition of cell division (Figure 16B). While colonies in the control group showed a relatively steady production of medusa buds throughout the experiment, animals treated with hydroxyurea exhibited a steep decrease in the production of early (stage 1) and intermediate (stage 2-4) buds (Figure 17F and 17G). By contrast, early buds that were present at the onset of development continued to develop up to 48 hours after treatment, as evident from the increase in the number of late stage buds (stage 5-7) (Figure 17H). The number of buds in stage 8 and above peaked around 48h after treatment and decreased until they were no longer observed after 4 days (Figure 17I). These results suggest that the initiation of medusae buds from and progression of early buds s require cell proliferation for their development to progress, however, later stages do not. The increase of buds from stage 5-7 and beyond stage 8, at 24h and 48h respectively, indicates that progression of the development occurred. However, lack of replacement of earlier stages of medusa buds and release of medusae are responsible for the decline in number of those stages after 48h.

**Figure 17:**
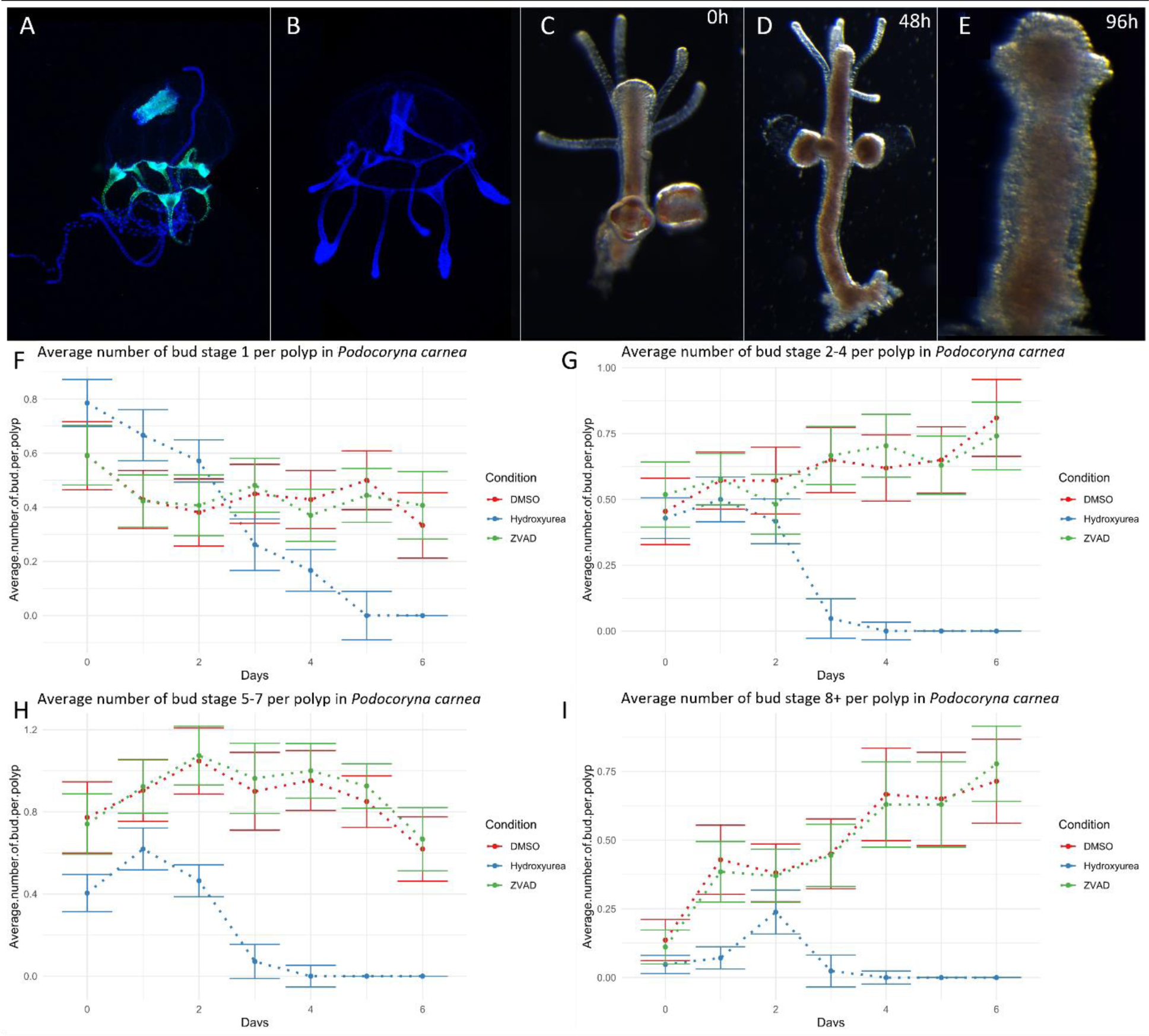
Effect of cell proliferation inhibition (hydroxyurea 10 µM) and cell apoptosis inhibition (Z-VAD-FMK 10 µM) on the development of *Podocoryna carnea*. Edu cell labeling on a newly released medusa incubated in 0.2% DMSO for 48h (A), green (Edu) and blue (Hoechst). Edu cell labeling on a newly released medusa incubated in 10µM hydroxyurea for 48h (B), green (Edu) and blue (Hoechst). Budding polyp treated with 10 µM hydroxyurea at 0h(C). Budding polyp treated with 10 µM hydroxyurea at 24h, showing regression of the buds (D). Budding polyp treated with 10 µM hydroxyurea at 48h, showing complete regression of the buds (E). Average number of early buds (stage 1) per polyp (F). Average number of intermediate buds (stage 2-4) per polyp (G). Average number of intermediate buds (stage 5-7) per polyp (H). Average number of late buds (stage 8+) per polyp (I).

The medusa of *Staurocladia* sp. is able to simultaneously bud multiple medusae at different stages of development. To address the role of cell proliferation in the development of the medusa, budding medusae of *Staurocladia* sp. were treated with 10 µM hydroxyurea, which effectively inhibited cell division (Figure 18B). Additionally, five successive generations of medusae were maintained in hydroxyurea. Throughout the experiment, the initial population of *Staurocladia* sp. treated with hydroxyurea did not show a significant (p=0.1122) difference in the number of medusa buds per medusa (Figure 18E). A significant difference in the number of buds per medusa was detected at the third generation of *Staurocladia* sp. maintained in hydroxyurea (Figure 18H).

**Figure 18:**
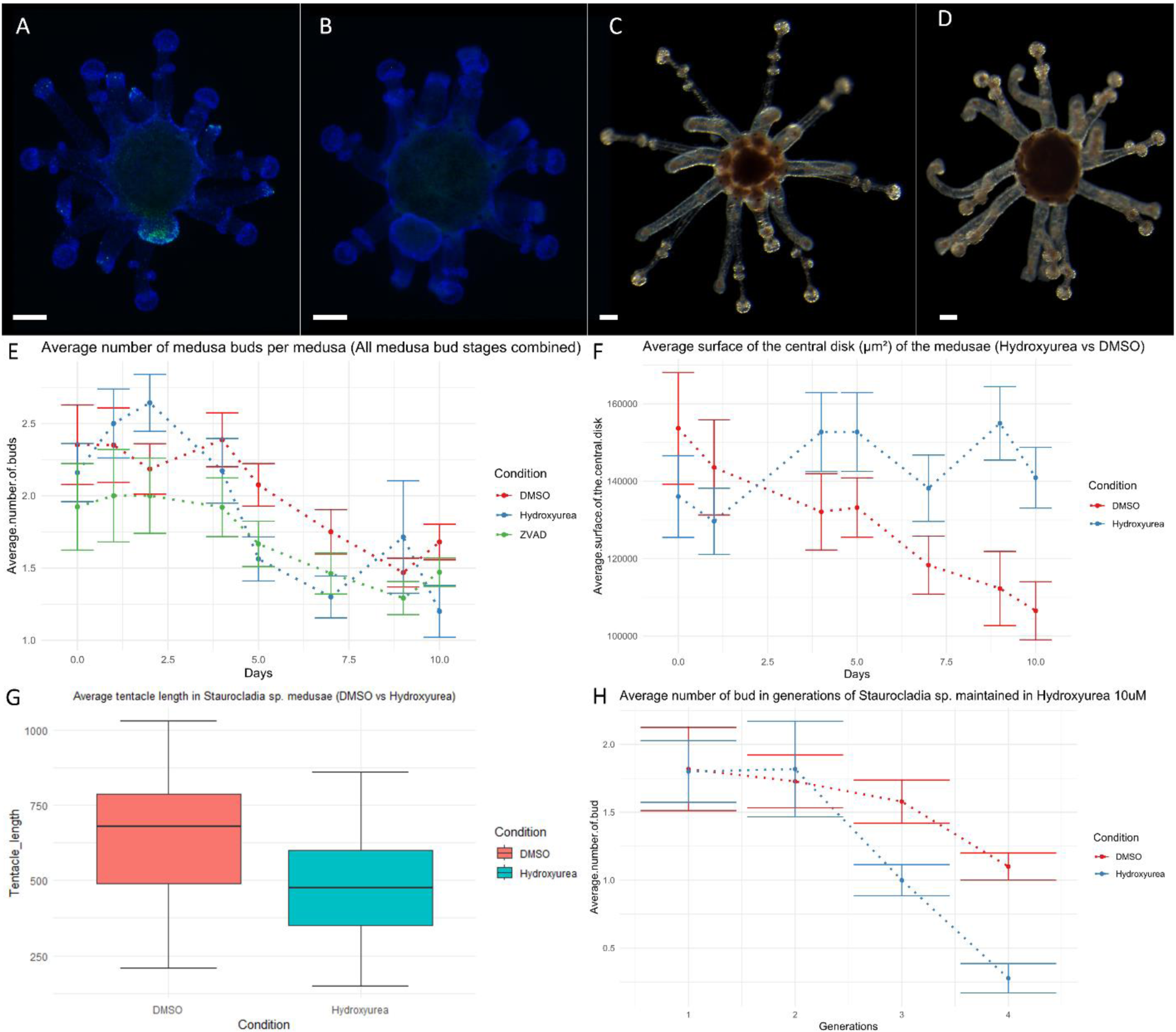
Effect of cell proliferation inhibition (hydroxyurea 10 µM) and cell apoptosis inhibition (Z-VAD-FMK 10 µM) on the development of *Staurocladia* sp. Edu cell labeling on an early budding medusa in 0.2% DMSO for 48h (A), green (Edu) and blue (Hoechst). Edu cell labeling on an early budding medusa incubated in 10µM hydroxyurea for 48h (B), green (Edu) and blue (Hoechst). Medusa after 10 days of incubation in DMSO 0.2% (C). Medusa after 10 days of incubation in 10 µM hydroxyurea (D). Average number of medusa buds per medusa (E). Average surface area of the central disk in medusae treated with hydroxyurea (F). Effect of hydroxyurea on the tentacle length after 10 days of treatment (G). Average number of medusa buds per medusa in successive generations of *Staurocladia* sp. (H).

As the medusae of *Staurocladia* sp. appeared able to bud multiple generations successively without cell division, we assessed whether medusa budding in the absence of cell division resulted in depletion of the pre-existing tissues. The surface area of the central disk was calculated by measuring the length (a) and the width (b) of the central disk. For the purpose of this study the central disk was considered to be an oval and the formula to determine the surface area was determined accordingly (abπ). Throughout the experiment, animals treated with hydroxyurea show a significantly larger surface area of the central disk (p=0.0023) (Figure 18F). After 10 days in hydroxyurea, the treated animals showed a significantly (p<0.0001) shorter tentacle length (Figure 18G), suggesting that cells from the tentacles are migrating to the central disk in the absence of cell proliferation.

In the generations of *Staurocladia* sp. maintained in hydroxyurea, the first generation did not show a significant difference in the average number of buds per medusa (p=0.9613), nor in the second generation (p=0.8236) (Figure 18H). At the third generation, a significantly lower number of buds per medusa was observed in the treated animals (p=0.0059), the effect was found greater in the fourth generation (p<0.0001, Figure 18H), presumably due to the depletion of cells.

## Discussion

Our comparative investigation provides insight into the role of cell proliferation, cell migration and programmed cell death in medusa development and in shaping medusa diversity. Previous studies have highlighted the role of cell proliferation in the adult hydromedusa of *Cladonema pacificum* (Fujita et al., 2019) (a close relative of *Staurocladia)* and found that cell proliferation in the exumbrella and tentacle bulbs play a role in body size (exumbrella growth) and tentacle morphogenesis. From the same study, the adult hydromedusa of *Cytaeis uchidae* and *Rathkea octopuntata* exhibit intense cell proliferation in the tentacle bulbs and the manubrium. Additionally, broad cell proliferation was detected in the early medusa bud of the medusa-budding-medusa of *R.octopunctata*. Our results are consistent with the cell proliferation patterns observed in this study, highlighting the conserved role of cell proliferation in medusa budding within Hydrozoa. Tentacle bulbs and the manubrium are known for their high cell turnover and their respective role in nematogenesis and gametogenesis. Upon release, the somatic development of the hydromedusa is often considered complete, and the conserved pattern of cell proliferation highlights conserved mechanisms in tissue and cell type maintenance.

The divergence of the development trajectories can however be observed in stages of the hydromedusa bud (Figure 19). The omnipresence of proliferating cells in the early bud coincides with the formation of the anatomically typical hydromedusae. Indeed, the *Hydractinia symbiolongicarpus* gonophore develops into a simple structure delimited by two single-layered epithelia (sporosac). This simplified gonophore architecture coincides with a weak and sparse cell proliferation pattern, compared to *P. carnea* and *B carcinicola* early buds, and more akin to baseline cell division. The eumedusoid of *B. carcinicola* represents an intermediate degree of the truncation of the gonophore development. The eumedusoid is short-lived and exhibits features of a fully developed medusa except for lacking tentacles, gonads, and a mouth. Interestingly, in the released medusoid of *B. carcinicola*, no cell proliferation is detected in the manubrium and the tentacle bulbs, the two areas of gonad and tentacle morphogenesis respectively. And in fact, the only apoptosis observed in hydrozoan medusae were in the short-lived released eumedusoids. Together these results suggest that the pattern of cell proliferation could be involved in the evolutionary heterochrony that gave rise to the diversification of hydractiniid life cycles. In this scenario, loss of cell proliferation in distally restricted or medusa specific structure could have resulted in simplification of the medusa in various lineages. Inhibition of cell proliferation in the late stages of the hydromedusa development does not result in loss of tentacles or gonads, indicating that either factors other than cell proliferation could account for the evolutionary loss of these structures or that development of these structures rely on cell proliferation occurring early during early stages.

**Figure 19:**
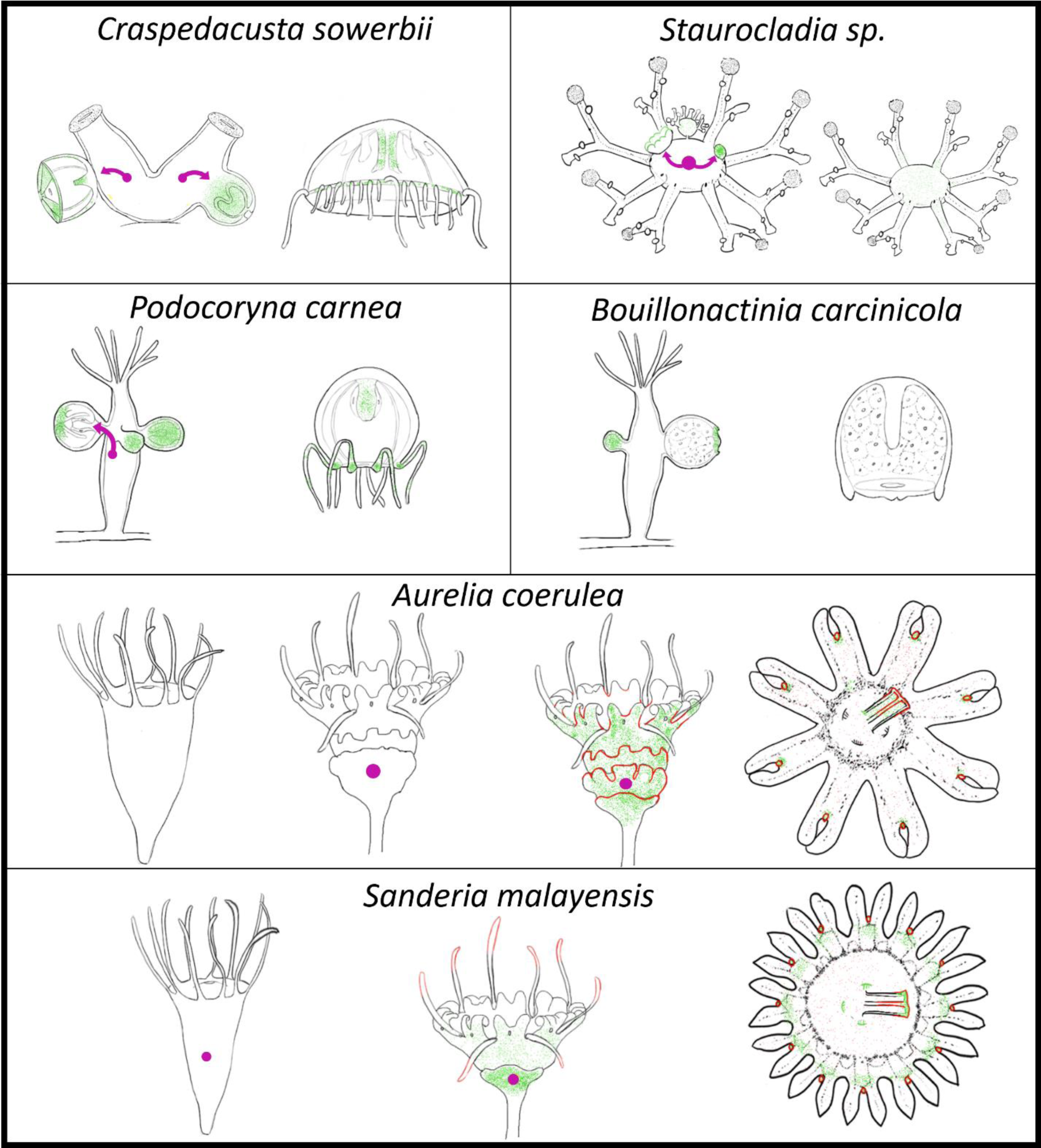
Summary of the pattern of cell proliferation, cell migration and apoptosis in study organisms. Cell proliferation (green), location of cells prior to migration (purple circle), migrating cells (purple arrow), apoptosis (red).

In the surveyed hydrozoan species, the timing of cell migration appears to follow that of cell proliferation. Indeed, in the two surveyed species possessing a fully developed hydromedusa derived from a polyp, migration of cells from the body column into the bud occurs at the late stages of development of the medusa bud (Figure 19). This could be explained by either recruitment of cells from the body column following cell proliferation or DiI signal dilution due to the extensive cell proliferation of migrated cells in the medusa bud. Interestingly, cell migration in *Staurocladia* sp. occurs very early during medusa development compared to the other surveyed hydrozoan species (Figure 19). The ability of *Staurocladia* sp. to complete development of the medusa in the absence of cell proliferation could be due to the early recruitment into the medusa bud of cells from the asexually budding medusa. Whether the extensive use of migrating cells in *Staurocladia* sp. is a conserved mechanism across medusa-to-medusa budding species remains to be investigated. By contrast, our results show that the development of the scyphomedusa does not recruit from a pre-existing pool of cells from the body column of the scyphistoma (Figure 19).

In the surveyed scyphozoan species, cell proliferation correlates with areas of active morphogenesis. Throughout the development of the scyphomedusa no proximo-distal gradient of cell proliferation was observed, however cell proliferation appears to show higher intensity in the epidermis early during strobilation and later on in the gastrodermis. This could indicate that, unlike hydrozoans, scyphozoan development relies on the successive morphogenesis of epidermal tissue and later in gastrodermal tissues. The rhopalium of the ephyra is a complex structure exhibiting various functionally specialized regions. When treating *A. coerulea* with a low dosage of hydroxyurea, strobilation and release of the ephyra was achieved. In this case, ephyrae lacked a pigment cup in the eye territory of the rhopalia. This is in contrast with *Staurocladia* sp. that maintains eye pigments in the absence of cell proliferation. Similar to scyphozoans, previous studies using Edu-tracing in cubozoan species *Tripedalia cystophora* and *Alatina moseri* show a specific pattern of cell proliferation in the rhopalia and the developing tentacles. More specifically, the cubozoan rhopalium exhibits cell proliferation in the pit and lens eyes (Gurska and Garm, 2014). Although no functional work was carried out of the cubozoan species, these results suggest that development of the eye in acraspedote medusae relies on a cell proliferation, despite being convergent between cubozoans and scyphozoans.

In contrast to hydrozoans the interplay between cell proliferation and cell migration from the body column of the polyp does not occur in the surveyed scyphozoans. During strobilation, cells from the body column labeled with DiI do not migrate to the developing ephyra and instead remain at the injection site (Figure 19). The disappearance of those cells in later stages of strobilation could be explained by intense cell proliferation, diluting the signal, or removal of those cells. Furthermore, labeling of cells present in the regressing tentacles of the early strobila indicates that those cells migrate into the rudiment, suggesting that those cells are excluded from the developing ephyra. It is worth noting these labeled cells were sporadically detected in the developing rhopalia in some disks.

In both hydrozoans and scyphozoans, apoptosis is involved in the metamorphosis of the planula and has been shown to be involved in the change in the nervous system architecture marking the transition from larva to polyp (Krasovec et al., 2021 and Yuan et al., 2008). However, the role of apoptosis during medusa development was not investigated prior to this study.

In contrast to hydrozoans, where apoptosis does not occur in medusa development), we found that scyphozoan medusa development is caspase dependent and likely relies on apoptosis. Throughout the development of the ephyra, apoptosis appears predominant in the epidermis, be it in the bell of the ephyra or the rhopalia (Figure 19). The first sign of cell apoptosis in the gastrodermis is detected in the ephyra within the manubrium (Figure 19). Throughout development cell proliferation and apoptosis were observed in the same territories suggesting that these processes co-occur.

Together our results show that the development of the scyphomedusa and the hydromedusa utilize vastly different cellular mechanisms. Although cell proliferation is required for the early development of both types of medusae, cell migration, and programmed cell death appear to play a distinct role in each medusa type. The crucial role of apoptosis during strobilation would indicate that its development relies on the restructuring of the polyp tissue. Stauromedusa and cubomedusae are known to develop from the transformation of the entire polyp (with few exceptions in cubozoans), however, the cellular mechanisms involved in their development are unknown. Nonetheless, medusa development through restructuring of polyp tissue, notably through extensive apoptosis and cell proliferation, would be inferred to be the ancestral condition in Acraspeda. In contrast, the hydromedusa development does not rely on apoptosis but instead on cell migration and cell proliferation. In all surveyed hydrozoan species, the medusa bud from the polyp consistently relies on restricted cell proliferation, in addition to cell migration into the bud (Figure 19), suggesting that those processes were ancestrally involved in the development of the hydromedusa in Hydrozoa. By contrast, medusa-to-medusa budding in *Staurocladia* does not appear to require cell division. The increase of the surface area of the central disk, in addition to the shrinkage of the tentacles of the medusa, and the ability of *Staurocladia* sp. to sustain medusa budding through several generations suggests that cells from the tentacles are contributing to the central disk tissue in the absence of cell division.

Our results provide evidence of the dissimilarity of the development between scyphozoan and hydrozoan medusae at the cellular level. These differences in cellular dynamics make the inference of ancestral mechanisms in the last common ancestor difficult. The ancestral medusa development likely would have consisted of structures derived from the polyp, utilizing cell proliferation. Later on, the divergence of cellular mechanisms could have shaped the divergence of the development in the two lineages. Alternatively, the dissimilarity of cellular dynamics could suggest a parallel evolution of the medusa form through independent developmental trajectories.

Investigation of the developmental toolkit orchestrating the striking differences in development between Hydrozoa and Acraspeda could further our understanding of the origin of the medusa form. Transcriptomic and gene expression pattern analyses of the life cycle of several medusozoan species have given some insights into the gene repertoire involved during medusa development. Comparative gene expression patterns between *Aurelia aurita* and *Clytia hemisphaerica* suggest changes in the expression pattern of conserved key developmental gene during transition from polyp to medusa is involved in medusa development (Kraus et al., 2015). However more recent studies have shown that several genes show a pattern of upregulation specifically during development and maintenance of the medusa stage (Khalturin et al., 2019, Leclere et al., 2019, Gold et al., 2019, Travert et al., 2023). Differential gene expression during medusa development points to the significant role of homeobox genes. Specifically, genes such as *Hox1, Drgx, Otx1, Msx* (Khalturin et al., 2019, Leclere et al., 2019, Sanders and Cartwright, 2015) show consistent medusa-specific expression across Scyphozoa and Hydrozoa. Additionally, the presence of the homeobox gene *Tlx* has been shown to strongly correlate with the presence of a medusa life cycle stage in cnidarians, with the independent losses of *Tlx* correlates with the multiple losses of the medusa stage (Travert et al., 2023). These homeobox genes implicated in g medusa development across medusozoans suggest a deep homology of the medusa at a molecular level, it remains unclear however, whether the structures wherein these genes are expressed can be homologized (e.g rhopalia and tentacle bulb). Our findings provide insight into the distinct cellular processes involved in medusae development and morphogenesis and demonstrates that an understanding of the complex evolutionary history of medusae requires comparisons at different levels of biological hierarchy.

## Methods

### Animal Care

*Podocoryna carnea* colonies were grown on microscope slides contained in slide racks and kept in artificial seawater (REEF CRYSTALS, Aquarium Systems) in a 7L Kreisel tank at room temperature (∼18°C) with a salinity of 29 ppt. Male and female colonies were kept in separate tanks. For the pharmacological treatments, pieces of colonies containing about 40 individuals of *Podocoryna carnea* male colonies (PCLH003) were transferred and grown for a month in polystyrene petri dishes (Cat# FB0875712) and kept in artificial seawater (REEF CRYSTALS, Aquarium Systems) at room temperature (∼18°C) with a salinity of 29 ppt. Colonies were fed two-day old Artemia nauplii twice a week. Prior to every experiment, *P. carnea* colonies were starved for four days.

*Podocoryna exigua* male colonies were collected in Roscoff, France, and were grown in the lab on the shell of a live *Tritia reticulata*. The colonies and the snail were and kept in artificial seawater (REEF CRYSTALS, Aquarium Systems) in a 1L glass Tupperware at room temperature (∼18°C) with a salinity of 29 ppt. Male and female colonies were kept in separate tanks. Colonies were fed two-day old Artemia nauplii twice a week. Prior to every experiment, *P. exigua* colonies were starved for four days and released budding polyps were collected. In *P. carnea*, removal of budding polyps from the colony and puncturing of the individual leads to regression of the early stages of the medusa bud. P.exigua budding polyps are spontaneously released from the colony, and puncturing of released budding individuals shows minimal to no impact on medusa bud development. P.exigua was selected as an alternative to *P. carnea* for the cell tracing experiment due to these properties. A partial sequence of the 16S rRNA was deposited on Genbank and can be accessed with the accession number OR644171.

Medusae of *Staurocladia sp* were obtained from a personal marine tank recently populated by multiple marine taxa collected by Gulf Specimen Marine Lab (Panacea, FL) including live rock samples. Animals were kept in artificial seawater (REEF CRYSTALS, Aquarium Systems) in 2L glass bowls at room temperature (∼18°C) with a salinity of 32 ppt. For the pharmacological treatments, medusae were transferred and in petri dishes (Cat# FB0875712) and kept in artificial seawater (REEF CRYSTALS, Aquarium Systems) at room temperature (∼18°C) with a salinity of 32 ppt. Medusae were fed two-day old Artemia nauplii twice a week. Prior to every experiment, *Staurocladia sp* were starved for four days. A partial sequence of the 16S rRNA was deposited on Genbank and can be accessed with the accession number OR644170.

*Aurelia coerulea* were grown on microscope slides contained in slide racks and kept in artificial seawater (REEF CRYSTALS, Aquarium Systems) in 2L glass bowls at room temperature (∼18°C) with a salinity of 29 ppt. Slides were transferred in glass bowls and strobilation was induced using 5-methoxy-2-methylindole at 250nM final concentration in artificial seawater (REEF CRYSTALS, Aquarium Systems) at room temperature (∼18°C) with a salinity of 29 ppt. Animals were kept in indole for 48h before the experiment and the medium was replaced with artificial sea water prior to each experiment. Polyps were fed two-day old Artemia nauplii twice a week. Prior to indole induction, *A. coerulea* were starved for four days.

*Sanderia malayensis* were grown on microscope slides contained in slide racks and kept in artificial seawater (REEF CRYSTALS, Aquarium Systems) in 2L glass bowls at room temperature (∼18°C) with a salinity of 29 ppt. Slides were transferred in glass bowls and strobilation was induced using 5-methoxy-2-methylindole at 250nM final concentration in artificial seawater (REEF CRYSTALS, Aquarium Systems) at room temperature (∼18°C) with a salinity of 29 ppt. Animals were kept in indole for 48h before the experiment and the medium was replaced with artificial sea water prior to each experiment. Polyps were fed two-day old Artemia nauplii twice a week. Prior to indole induction, *S. malayensis* starved for four days. A partial sequence of the 16S rRNA was deposited on Genbank and can be accessed with the accession number OR644169.

*Bouillonactinia carcinicola* female colonies were collected on the gastropod shell inhabited by *Clibanarius vittatus* ordered from Gulf Specimen Marine Lab (Panacea, FL). The colonies and the crab were kept in artificial seawater (REEF CRYSTALS, Aquarium Systems) in a 1L glass dish at room temperature (∼18°C) with a salinity of 29 ppt. Colonies were fed two-day old *Artemia* nauplii twice a week. Colonies used for staining were grown on pieces (1-2cm) of the gastropod shell. A partial sequence of the 16S rRNA was deposited on Genbank and can be accessed with the accession number OR644168.

*Hydractinia symbiolongicarpus* colonies (strain 296-10) were grown on microscope slides contained in slide racks and kept in artificial seawater (Red Sea Coral Pro Salt) in a 1L plastic dish at room temperature (∼18°C) with a salinity of 29 ppt. Pieces of colonies containing about 20 individuals of female *H. symbiolongicarpus* were transferred in 24-well plates in artificial seawater (Red Sea Coral Pro Salt) at room temperature (∼18°C) with a salinity of 29 ppt. Colonies were fed two-day old Artemia nauplii twice a week. Prior to every experiment, *H. symbiolongicarpus* colonies were starved for four days.

Polyps of *Craspedacusta sowerbii* (strain from Japan) were grown in 150mm tissue culture dish (MIDSCI, TP93150) and kept in Hydra medium (0.03mM KNO3, 1.0mM CaCl2.2H2O, 0.1mM MgCl2.6H2O, 0.5mM NaHCO3, 0.08mM MgSO4.7H2O). For the pharmacological treatments, budding polyps were transferred to polystyrene petri dishes (Cat# FB0875712) prior to every experiment. *Craspedacusta sowerbii* polyps were starved a week prior to any experiment.

### Cell labeling

#### Edu-labeling

Animals were incubated 20 μM 5-ethynyl-2′-deoxyuridine (Click-iT™ EdU Cell Proliferation Kit for Imaging, Alexa Fluor™ 488 dye, Cat# C10337) in 2ml of their respective artificial medium in 24 wells plates. The animals were relaxed with menthol (1mg/ml) for 30min except for *Craspedacusta sowerbii* which was not relaxed. The medium was replaced, and animals were fixed in two steps. The animals were fixed in 4% PFA in fresh filtered (using 0.22 µm Millex® PVDF syringe filter) medium for 30min, followed by a 30min fixation in 4%PFA-PBS. The samples were washed 3 times in 0.3% Triton-X100 -PBS for 15min with rocking. The samples were incubated with the EdU reaction cocktail (according to the manufacturer protocol, Cat# C10337) for 30 min in the dark. The samples were washed 3 times in 0.3% Triton-X100 -PBS for 15min with rocking, the first wash was added with Hoechst (1:500). Samples were placed in 90% glycerol and incubated overnight at 4C with rocking. Samples were then imaged.

#### TUNEL assay

The animals were relaxed with menthol (1mg/ml) for 30min except for *Craspedacusta sowerbii* which was not relaxed. The medium was replaced, and animals were fixed in two steps. The animals were fixed in 4% PFA in fresh filtered (using 0.22 µm Millex® PVDF syringe filter) medium for 30min, followed by a 30min fixation in 4%PFA-PBS. The samples were washed twice in 0.3% Triton-X100-PBS for 15min with rocking. The samples were incubated proteinase K (provided in Click-iT™ Plus TUNEL Assay Kits for In Situ Apoptosis Detection, Cat# C10617) for 30min, then washed twice in 0.3%PBS-Triton for 15 minutes. Samples were incubated in 4% PFA-PBS with rocking, then washed twice in 0.3%PBS-Triton for 15 minutes. Samples were washed with DEPC treated water for 3min with rocking. The TdT reaction and Click-iT plus reaction were carried out according to the manufacturer protocol (Cat# C10618). Samples were washed in 3% BSA-PBS for 5 min, then rinsed in PBS for 3 min. The samples were washed 3 times in 0.3% Triton-X100 -PBS for 15min with rocking, the first wash was added with Hoechst (1:500). Samples were placed in 90% glycerol and incubated overnight at 4C with rocking. Samples were then imaged.

#### Cell-tracing

The Dil Stain (1,1’-Dioctadecyl-3,3,3’,3’-Tetramethylindocarbocyanine Perchlorate (’DiI’; DiIC18(3))) working solution was dissolving DiI in 100% DMSO (Cat# D282, for a final concentration of 2mM). The animals were relaxed with menthol (1mg/ml) for 30min except *Craspedacusta sowerbii* which was not relaxed. About 20-30nl of the DiI working solution was pressured injected into the mesoglea of the animals. For the scyphozoan species (*S. malayensis* and *A. coerulea*), polyps were induced with indole 48h prior to the injection. Scyphostomae were injected in the lower portion of the body column, budding polyps of *P.exigua* and *C. sowerbii* were injected below the budding zone. Budding medusae of *Staurocladia sp* were injected in the middle of the exumbrella. After injection the animals were washed in fresh medium with rocking for 2h to remove the surplus DiI, then were imaged. When possible, the animals were imaged in bright field, otherwise labeled cells were imaged through epifluorescence microscopy.

### Pharmacological treatments

#### Hydroxyurea

Animals were incubated in 10 µM Hydroxyurea (Cat# A10831.03, dissolved in 100% DMSO) in their respective fresh filtered (using 0.22 µm Millex® PVDF syringe filter) medium. The medium was renewed every 24h. For *A. coerulea* an additional experiment was carried out using 1 µM Hydroxyurea. Incubation in DMSO was used as a control, the concentration of DMSO for the control group was determined according to the highest final DMSO concentration used among all pharmacological treatments (0.5% for the scyphozoan species and 0.2% for the hydrozoan species). The progression of the medusa development was recorded every 24h.

#### Pan-caspase inhibitor Z-VAD-FMK

Animals were incubated in Z-VAD-FMK (Promega, Cat# G7232) at a final concentration of 10 µM for the hydrozoan species and 100 µM for the scyphozoan species in their respective fresh filtered (using 0.22 µm Millex® PVDF syringe filter) medium. he medium was renewed every 24h. Incubation in DMSO was used as a control, the concentration of DMSO for the control group was determined according to the highest final DMSO concentration used among all pharmacological treatments (0.5% for the scyphozoan species and 0.2% for the hydrozoan species). The progression of the medusa development was recorded every 24h. Initially, animals were treated with 10µMZ-VAD-FMK, and no significant effect was detected in any of surveyed species (p-value greater than 0.59 for all species tested). The use of 10µM of Z-VAD-FMK did not result in a decrease of apoptotic bodies detected through TUNEL assay in the scyphozoan species. A decrease in the number of apoptotic bodies was detected in the scyphozoan species using 100µM Z-VAD-FMK, thus that dosage was used to assess the role of apoptosis in the two scyphozoan species. Hydrozoan species did not exhibit any signs of apoptosis as revealed by the TUNEL assay (Figure S1); thus, 10µM Z-VAD-FMK was used, consistent with the dosage used in the hydroxyurea experiments and previous study showing that 10 µM effectively inhibit caspases in *Hydractinia* (Seipp et al., 2006).

#### Statistical analyses

Statistical analyses were performed in RStudio v1.3.1093. Differences between the means were considered significant when p < 0.05, using unpaired two-tailed Student’s t-tests or Welch two-sample t-tests. Line charts were generated in R using ggplot.

#### Microscopy and imaging

Specimens were imaged on either a Leica DM5000 B compound microscope with a Lumenera INFINITY 3s camera or a Leica MZ16F with a Lumenera INFINITY 5. Images from different channels were merged using ImageJ v1.53e. Images were adjusted for brightness and contrast; any adjustments were applied to the entire image, not parts. DiI is excited at 550nm and emits at 594nm (red), DiI labeling was colorized in yellow using ImageJ v1.53e on Figures 7, 8 and 9 for clarity. Brightness was adjusted on fluorescent images (Photoshop vs. 24.4.3) using the brightness setting on the entire image.

## Supplementary Figures

**Figure S1:**
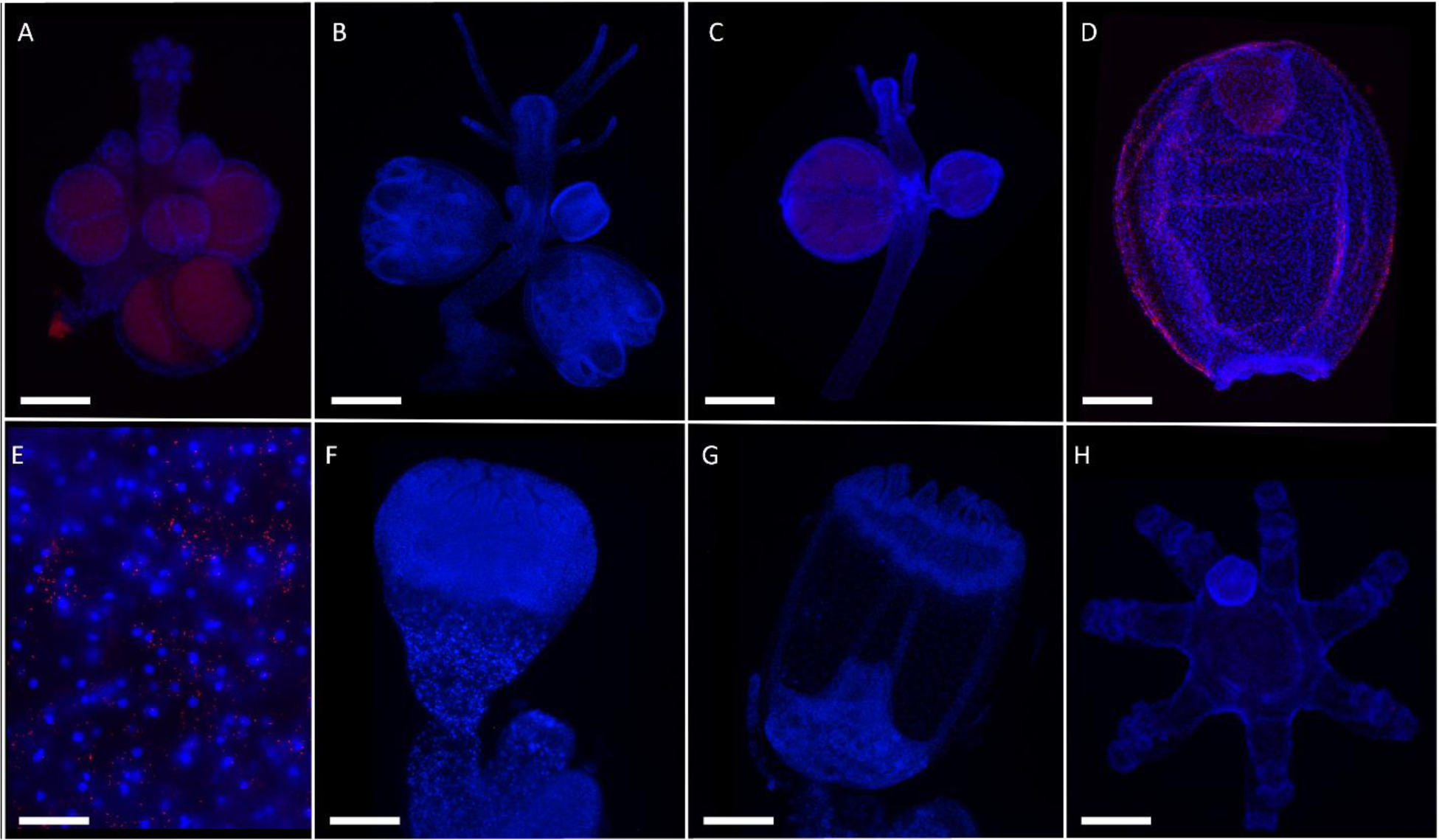
TUNEL assay in *Hydractinia symbiolongicarpus* (A), *Podocoryna carnea* (B), *Bouillonactinia carcinicola* (C-E), *Craspedacusta sowerbi*i (F-G), *Staurocladia* sp. (H). Gonozoid (A), budding polyp (B-C), eumedusoid (D), exumbrella of the medusoid (E), intermediate medusa bud (F), late medusa bud (G), budding medusa (H). Scale bars: 100 μm (A-C), 200 μm (D, F, G, H), 10μm (E). Oocytes in A and C exhibit autofluorescence and the red signal observed does not correspond to TUNEL positive signal.

**Figure S2:**
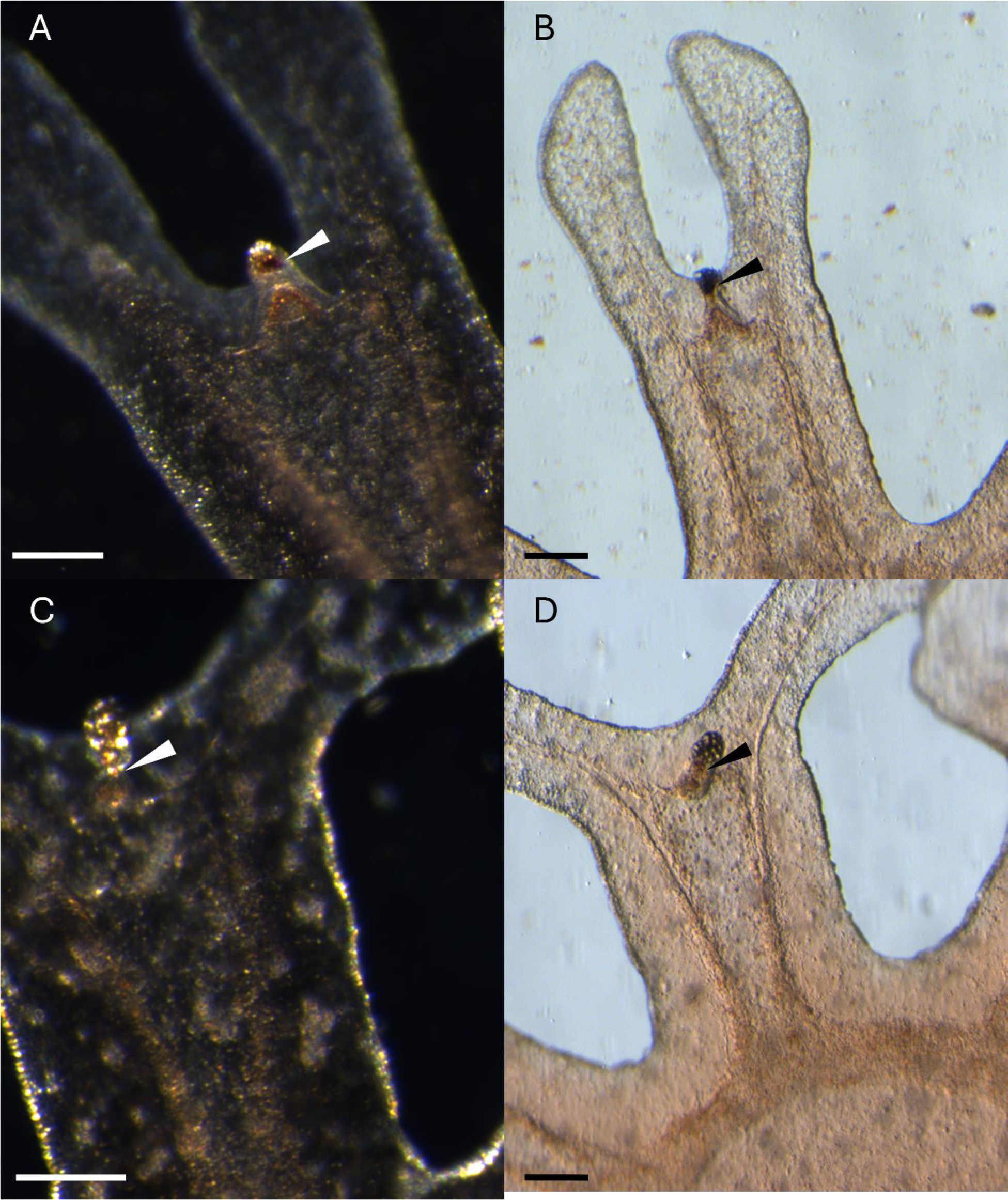
Rhopalium morphology of *Aurelia coerulea* ephyrae released from strobilae treated with 1μM hydroxyurea. Animals treated with 0.5% DMSO for 168h (A-B). Animals treated with 1μM hydroxyurea for 168h (C-D). The arrowhead indicates the area of the rhopalium bearing the pigment spot. Scale bars: 150μm.

## References

1. Petersen KW 1979. Development of coloniality in Hydrozoa. In: Biology and Systematics of Colonial Animals. G Larwood & BR Rosen, eds., pp. 105–139. Academic Press, London.

2. Bridge, D., Cunningham, C. W., DeSalle, R., & Buss, L. W. (1995). Class-level relationships in the phylum Cnidaria: Molecular and morphological evidence. Molecular Biology and Evolution, 12(4), 679–689.

3. Zapata, F., Goetz, F. E., Smith, S. A., Howison, M., Siebert, S., Church, S. H., Sanders, S. M., Ames, C. L., McFadden, C. S., France, S. C., Daly, M., Collins, A. G., Haddock, S. H. D., Dunn, C. W., & Cartwright, P. (2015). Phylogenomic Analyses Support Traditional Relationships within Cnidaria. PLOS ONE, 10(10), e0139068.

4. Collins, A. G., Schuchert, P., Marques, A. C., Jankowski, T., Medina, M., & Schierwater, B. (2006). Medusozoan phylogeny and character evolution clarified by new large and small subunit rDNA data and an assessment of the utility of phylogenetic mixture models. Systematic biology, 55(1), 97–115.

5. Simion, P., Philippe, H., Baurain, D., Jager, M., Richter, D. J., Di Franco, A., Roure, B., Satoh, N., Quéinnec, É., Ereskovsky, A., Lapébie, P., Corre, E., Delsuc, F., King, N., Wörheide, G., & Manuel, M. (2017). A Large and Consistent Phylogenomic Dataset Supports Sponges as the Sister Group to All Other Animals. Current Biology, 27(7), 958–967.

6. Gegenbaur, C. (1856). Versuch eines Systemes der Medusen, mit Beschreibung neuer oder wenig gekannter Formen; zugleich ein Beitrag zur Kenntniss der Fauna des Mittelmeeres. Wilhelm Engelmann.

7. Jarms, G., Båmstedt, U., Tiemann, H., Martinussen, M. B., Fosså, J. H., & Høisœter, T. (1999). The holopelagic life cycle of the deep-sea medusa Periphylla periphylla (Scyphozoa, Coronatae). Sarsia, 84(1), 55–65.

8. Petersen, K. W. (1990). Evolution and taxonomy in capitate hydroids and medusae (Cnidaria: Hydrozoa). Zoological Journal of the Linnean Society, 100(2), 101–231.

9. Bentlage, B., Osborn, K. J., Lindsay, D. J., Hopcroft, R. R., Raskoff, K. A., & Collins, A. G. (2018). Loss of metagenesis and evolution of a parasitic life style in a group of open-ocean jellyfish. Molecular Phylogenetics and Evolution, 124, 50–59.

10. Cartwright, P., & Nawrocki, A. M. (2010). Character Evolution in Hydrozoa (phylum Cnidaria). Integrative and Comparative Biology, 50(3), 456–472.

11. Leclère, L., Schuchert, P., Cruaud, C., Couloux, A., & Manuel, M. (2009). Molecular Phylogenetics of Thecata (Hydrozoa, Cnidaria) Reveals Long-Term Maintenance of Life History Traits despite High Frequency of Recent Character Changes. Systematic Biology, 58(5), 509–526.

12. Boelsterli, U. (1977). An electron microscopic study of early developmental stages, myogenesis, oogenesis and cnidogenesis in the anthomedusa, Podocoryne carnea M. Sars. Journal of Morphology, 154(2), 259–289.

13. Lesh-Laurie GE & Suchy PE 1991. Cnidaria: Scyphozoa and Cubozoa. In: Microscopic Anatomy of the Invertebrates, Vol. 2. Harrison FW & Westfall JA, eds., pp. 185–266. Wiley-Liss, New York.

14. Seipel, K., & Schmid, V. (2005). Evolution of striated muscle: Jellyfish and the origin of triploblasty. Developmental Biology, 282(1), 14–26.

15. Satterlie, R. A. (2008). Control of swimming in the hydrozoan jellyfish Aequorea victoria: subumbrellar organization and local inhibition. Journal of experimental Biology, 211(21), 3467–3477.

16. CarréC&CarréD1994. Ordre des siphonophores. In: Traité de Zoologie, Tome III, Fascicule 2— Cnidaires et Ctenaires.

17. Kühn, A. (1910). Die Entwicklung der Geschlechtsindividuen der Hydromedusen. Aus dem Zoologischen Institut der Universität Freiburg i. B.

18. Thomas MB & Edwards NC 1991. Cnidaria: Hydrozoa. In: Microscopic Anatomy of the Invertebrates, Vol. 2. Harrison FW & Westfall JA, eds., pp. 91–183. Wiley-Liss, New York.

19. Koizumi, O., Hamada, S., Minobe, S., Hamaguchi-Hamada, K., Kurumata-Shigeto, M., Nakamura, M., & Namikawa, H. (2015). The nerve ring in cnidarians: Its presence and structure in hydrozoan medusae. Zoology, 118(2), 79–88.

20. Gröger, H., & Schmid, V. (2000). Nerve net differentiation in medusa development of *Podocoryne carnea*. Scientia Marina, 64(S1), Article S1.

21. Horridge, G. A. (1956). The Nerves and Muscles of Medusae: V. Double Innervation in Scyphozoa. Journal of Experimental Biology, 33(2), 366–383.

22. Nakanishi, N., Hartenstein, V., & Jacobs, D. K. (2009). Development of the rhopalial nervous system in Aurelia sp.1 (Cnidaria, Scyphozoa). Development Genes and Evolution, 219(6), 301–317.

23. Westlake, H. E., & Page, L. R. (2017). Muscle and nerve net organization in stalked jellyfish (Medusozoa: Staurozoa). Journal of Morphology, 278(1), 29–49.

24. Garm, A., Ekström, P., Boudes, M., & Nilsson, D.-E. (2006). Rhopalia are integrated parts of the central nervous system in box jellyfish. Cell and Tissue Research, 325(2), 333–343.

25. Nielsen, S. K. D., Koch, T. L., Wiisbye, S. H., Grimmelikhuijzen, C. J. P., & Garm, A. (2021). Neuropeptide expression in the box jellyfish *Tripedalia cystophora*—New insights into the complexity of a “simple” nervous system. Journal of Comparative Neurology, 529(11), 2865–2882.

26. Picciani, N., Kerlin, J. R., Sierra, N., Swafford, A. J. M., Ramirez, M. D., Roberts, N. G., Cannon, J. T., Daly, M., & Oakley, T. H. (2018). Prolific Origination of Eyes in Cnidaria with Co-option of Non-visual Opsins. Current Biology, 28(15), 2413–2419.e4.

27. Bouillon J., 1985 - Essai de classification des Hydropolypes-Hydromeduses (Hydrozoa-Cnidaria). Indo-Mal. Zool., 1 : 29-243.

28. Bouillon J. (1994). Classe des Hydrozoaires, in Traité de Zoologie (Eds P. P. Grasse, D. Doumenc), Masson, Paris, 3,29-416

29. Mayer, A.G. (1910) Medusae of the world. Volume I. The hydromedusae. Carnegie Institution of Washington, Publication, 109, 1–230

30. Chapman, DM. (1985). X-ray microanalysis of selected coelenterate statoliths. J. Mar. Biol. Assoc. U.K. 65: 617–627.

31. Bouillon J. (1978). Hydroméduses de la Mer de Bismarck (Papouasie Nouvelle Guinée). Partie II: Limnomedusa, Narcomedusa, Trachymedusa et Laingiomedusa (sous-classe nov.). Cah. Biol. Mar. 19: 473–483.

32. Thiel H 1966. The evolution of the Scyphozoa, a review. In: Cnidaria and their Evolution. Rees WJ, ed., pp. 77–117. Academic Press, London.

33. Adonin, L. S., Shaposhnikova, T. G., & Podgornaya, O. (2012). Aurelia aurita (Cnidaria) Oocytes’ Contact Plate Structure and Development. PLOS ONE, 7(11), e46542.

34. Garcia-Rodriguez, J., Lewis Ames, C., Marian, J., & Marques, A. (2018). Gonadal histology of box jellyfish (Cnidaria: Cubozoa) reveals variation between internal fertilizing species Alatina alata (Alatinidae) and Copula sivickisi (Tripedaliidae). Journal of Morphology, 279.

35. Berrill, M. (1963). Comparative functional morphology of the Stauromedusae. Canadian Journal of Zoology, 41(5), 741–752.

36. Bouillon, Gravili, C., F, P., Gili, J.-M., & Boero, F. (2006). An Introduction to Hydrozoa. Mémoires Du Muséum National d’Histoire Naturelle, 194, 1–591.

37. 1994b. Classe des hydrozoaires (Hydrozoa Owen, 1843). In: Traité deZoologie, Tome III, Fascicule 2—Cnidaires et Ctenaires. Bouillon J, Carré C,Carré D,Franc A, Goy J, Hernandez-Nicaise M-L, Tiffon Y, van de Vyver D, & Wade M, eds., pp. 29–416. Masson, Paris.

38. Berrill, N. J. (1950). Development and medusa-bud formation in the Hydromedusae. The quarterly review of Biology, 25(3), 292–316.

39. Seipel, K., Yanze, N., & Schmid, V. (2004). The germ line and somatic stem cell gene Cniwi in the jellyfish Podocoryne carnea. The International Journal of Developmental Biology, 48(1), 1–7.

40. Chari, T., Weissbourd, B., Gehring, J., Ferraioli, A., Leclère, L., Herl, M., Gao, F., Chevalier, S., Copley, R. R., Houliston, E., Anderson, D. J., & Pachter, L. (2021). Whole-animal multiplexed single-cell RNA-seq reveals transcriptional shifts across Clytia medusa cell types. Science Advances, 7(48), eabh1683.

41. Marques, A. C., & Collins, A. G. (2004). Cladistic analysis of Medusozoa and cnidarian evolution. Invertebrate Biology, 123(1), 23–42.

42. Helm, R. R. (2018). Evolution and development of scyphozoan jellyfish. Biological Reviews, 93(2), 1228–1250.

43. Helm, R. R., Tiozzo, S., Lilley, M. K. S., Lombard, F., & Dunn, C. W. (2015). Comparative muscle development of scyphozoan jellyfish with simple and complex life cycles. EvoDevo, 6(1), 11.

44. Schwab, W. E. (1977). The ontogeny of swimming behavior in the scyphozoan, Aurelia aurita. I. Electrophysiological analysis. The Biological Bulletin, 152(2), 233–250.

45. Sandrini, L. R., & Avian, M. (1983). Biological cycle of Pelagia noctiluca: Morphological aspects of the development from planula to ephyra. Marine Biology, 74(2), 169–174.

46. Kikinger, R., & Salvini-Plawen, L. von. (1995). Development From Polyp to Stauromedusa in Stylocoronella (Cnidaria: Scyphozoa). Journal of the Marine Biological Association of the United Kingdom, 75(4), 899–912.

47. Stangl K, Salvini-Plawen Lv, & Holstein TW 2002. Staging and induction of medusa metamorphosis in Carybdea marsupialis (Cnidaria, Cubozoa). Vie Milieu 52: 131–140.

48. Toshino, S., Miyake, H., Ohtsuka, S., Adachi, A., Kondo, Y., Okada, S., Hirabayashi, T., & Hiratsuka, T. (2015). Monodisc strobilation in Japanese giant box jellyfish Morbakka virulenta (Kishinouye, 1910): A strong implication of phylogenetic similarity between Cubozoa and Scyphozoa. Evolution & Development, 17(4), 231–239.

49. Straehler-Pohl, I., & Jarms, G. (2005). Life cycle of Carybdea marsupialis Linnaeus, 1758 (Cubozoa, Carybdeidae) reveals metamorphosis to be a modified strobilation. Marine Biology, 147(6), 1271–1277.

50. Berrill, N. J. (1949). Form and growth in the development of a scyphomedusa. The Biological Bulletin, 96(3), 283–292.

51. Helm, R. R., & Dunn, C. W. (2017). Indoles induce metamorphosis in a broad diversity of jellyfish, but not in a crown jelly (Coronatae). PLOS ONE, 12(12), e0188601.

52. Fujita, S., Kuranaga, E., & Nakajima, Y. (2019). Cell proliferation controls body size growth, tentacle morphogenesis, and regeneration in hydrozoan jellyfish Cladonema pacificum. PeerJ, 7, e7579.

53. Gurska, D., & Garm, A. (2014). Cell Proliferation in Cubozoan Jellyfish Tripedalia cystophora and Alatina moseri. PLoS ONE, 9(7), e102628.

54. Krasovec, G., Pottin, K., Rosello, M., Quéinnec, É., & Chambon, J.-P. (2021). Apoptosis and cell proliferation during metamorphosis of the planula larva of Clytia hemisphaerica (Hydrozoa, Cnidaria). Developmental Dynamics, 250(12), 1739–1758.

55. Yuan, D., Nakanishi, N., Jacobs, D. K., & Hartenstein, V. (2008). Embryonic development and metamorphosis of the scyphozoan Aurelia. Development Genes and Evolution, 218(10), 525–539.

56. Kraus, J. E. M., Fredman, D., Wang, W., Khalturin, K., & Technau, U. (2015). Adoption of conserved developmental genes in development and origin of the medusa body plan. EvoDevo, 6(1), 23.

57. Khalturin, K., Shinzato, C., Khalturina, M., Hamada, M., Fujie, M., Koyanagi, R., Kanda, M., Goto, H., Anton-Erxleben, F., Toyokawa, M., Toshino, S., & Satoh, N. (2019). Medusozoan genomes inform the evolution of the jellyfish body plan. Nature Ecology & Evolution, 3(5), 811–822.

58. Leclère, L., Horin, C., Chevalier, S., Lapébie, P., Dru, P., Peron, S., Jager, M., Condamine, T., Pottin, K., Romano, S., Steger, J., Sinigaglia, C., Barreau, C., Quiroga Artigas, G., Ruggiero, A., Fourrage, C., Kraus, J. E. M., Poulain, J., Aury, J.-M., … Copley, R. R. (2019). The genome of the jellyfish Clytia hemisphaerica and the evolution of the cnidarian life-cycle. Nature Ecology & Evolution, 3(5), 801–810.

59. Gold, D. A., Katsuki, T., Li, Y., Yan, X., Regulski, M., Ibberson, D., Holstein, T., Steele, R. E., Jacobs, D. K., & Greenspan, R. J. (2019). The genome of the jellyfish Aurelia and the evolution of animal complexity. Nature Ecology & Evolution, 3(1), 96–104.

60. Travert, M., Boohar, R., Sanders, S. M., Boosten, M., Leclère, L., Steele, R. E., & Cartwright, P. (2023). Coevolution of the Tlx homeobox gene with medusa development (Cnidaria: Medusozoa). Communications Biology, 6(1), 1–12.

61. Sanders, S. M., & Cartwright, P. (2015). Interspecific Differential Expression Analysis of RNA-Seq Data Yields Insight into Life Cycle Variation in Hydractiniid Hydrozoans. Genome Biology and Evolution, 7(8), 2417–2431.

62. Seipp, S., Wittig, K., Stiening, B., Bottger, A., & Leitz, T. (2005). Metamorphosis of Hydractinia echinata (Cnidaria) is caspase-dependent. The International journal of developmental biology, 50(1), 63–70.

