## Supplementary figures and images for "Comparative investigations of cellular dynamics in the development of medusae (Cnidaria: Medusozoa)"

### Figure S1

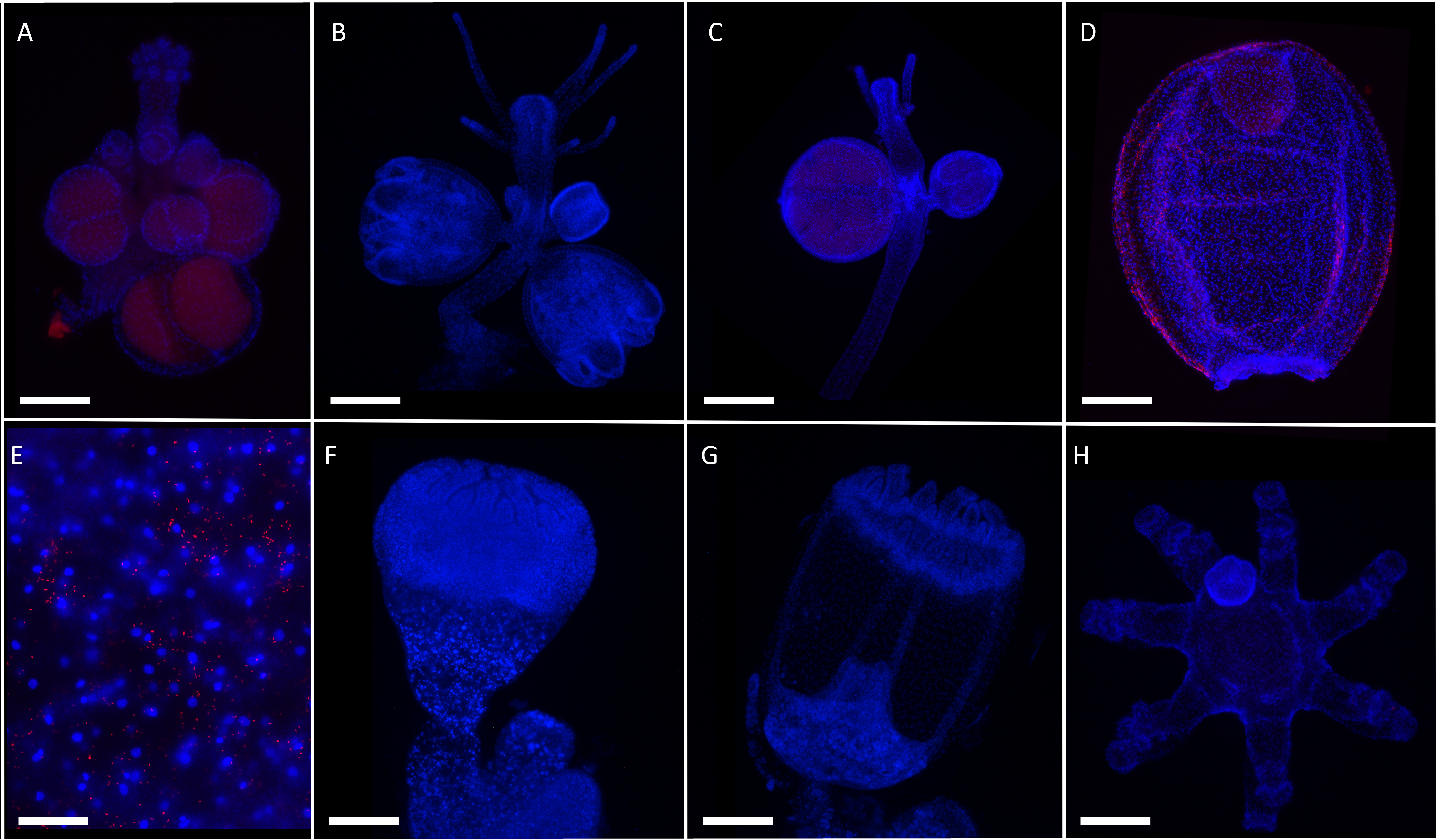

### Figure S2

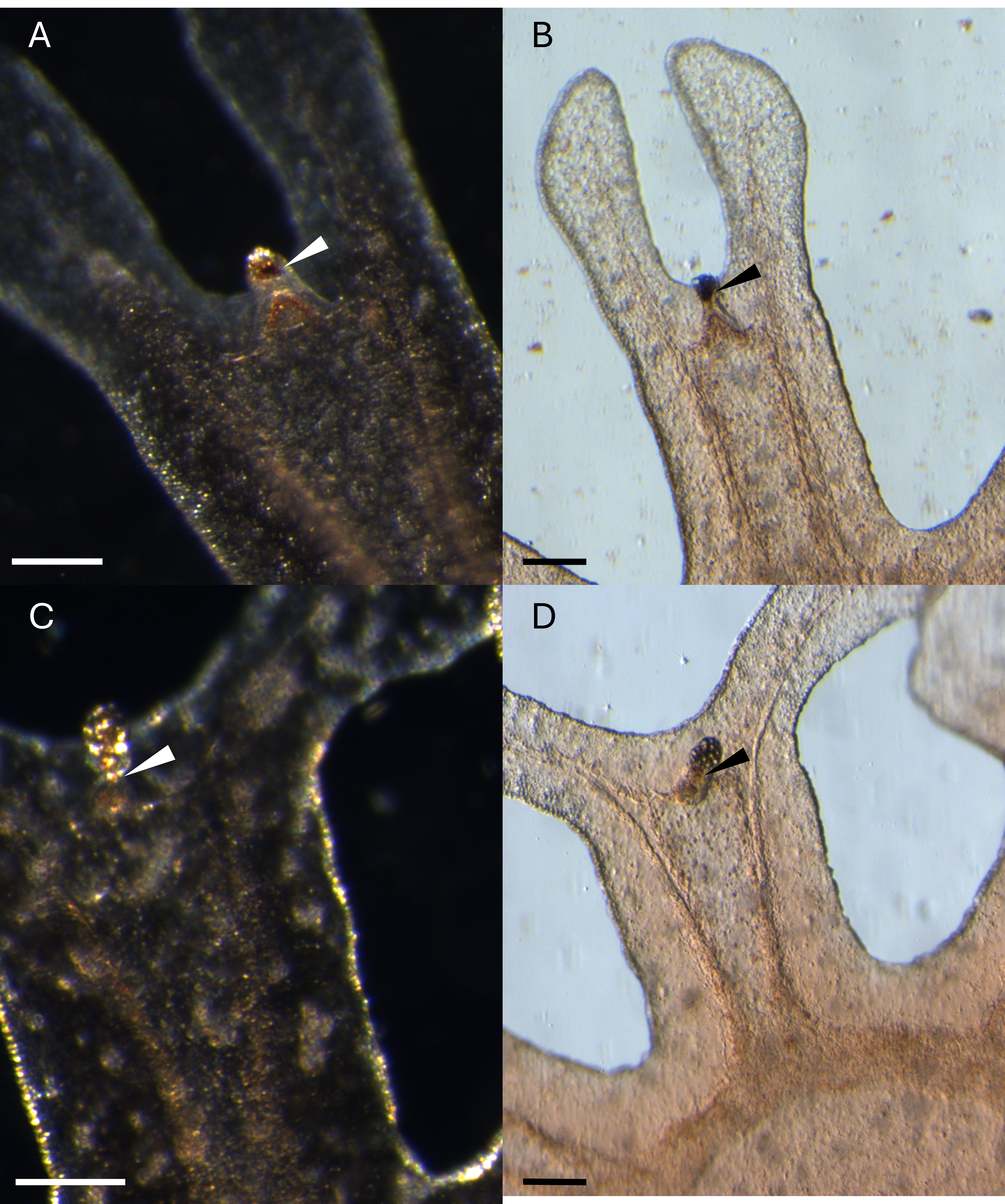
